# Cytokinin-mediated repression of jacalin lectins reinforces root immunity

**DOI:** 10.64898/2026.05.15.725414

**Authors:** Abin Panackal George, Varsha Koolath, Krishna Kodappully Das, Arpita Das, Subiya Haque, Irfana Thesni Karuvanthodi, Kiran Ambatipudi, Santosh B Satbhai, Eswarayya Ramireddy

**Author notes:** **Corresponding author details:** Dr. Eswarayya Ramireddy, Department of Biology, Indian Institute of Science Education and Research (IISER) Tirupati, Tirupati 517619, Andhra Pradesh, India.

## Abstract

The root cap is essential for perceiving environmental cues surrounding the root. However, the molecular mechanisms underlying root cap-mediated immunity and how it defends against invading pathogens remain largely unresolved.

Our results indicate that cytokinin plays a major role in regulating soil-borne pathogen such as *Ralstonia pseudosolanacearum* load around the root and root cap. As *Ralstonia* populations increase, cytokinin signalling is activated and represses the expression of its downstream signalling targets such as root cap-specific proteins *JAL10* and *JAL20*, to impart the tolerance against the *Ralstonia*.

The functional analysis jacalin-associated lectin family proteins *JAL10* and *JAL20*, revealed that *loss-of-function* leads to enhance tolerance to *Ralstonia* whereas *gain-of-function* leads to susceptibility compared to Col-0. Our Glycoproteomic and metabolomic analyses indicate that *JAL10* and *JAL20* act as negative regulators of cell wall remodelling and likely to promotes cell wall thickening, thereby enhancing resistance to soil-borne infections.

The knockdown of ortholog of JAL protein in Tomato also revealed its conserved function in imparting tolerance to *Ralstonia pseudosolanacearum.* Further we also show downregulation of JALs by other soil-borne pathogen infection, suggesting that cytokinin might protecting the vulnerable areas of root tip regions by regulating the expression of root cap-specific JALs and thereby fortifying the cell wall.

## Introduction

Roots grow in soil environments that are highly diverse and constantly changing. From the earliest stages of development, the emerging radicle can sense its surroundings and initiate adaptive responses suited to the prevailing environment. At the time of emergence, the radicle tip is initially covered by a thin cuticular layer that provides protection during early growth. Disruption or premature removal of the cuticle can lead to hypersensitivity and may compromise seedling viability. As development progresses, the cuticle protective layer is replaced by the root cap, a specialized structure that facilitates soil penetration and serves as a critical sensory interface (Berhin et al., 2019). As the root advances through the soil, the root cap plays a central role in environmental perception (Kumpf & Nowack, 2015; Ganesh et al., 2022). Root-cap integrates multiple external cues, including gravity (Tsugeki et al., 1999), water availability (Miyazawa, & Takahashi, 2020), nutrient gradients (Kanno et al., 2016), and the presence of potentially toxic compounds (Ryan, et al 1993). This sensory capacity enables the plant to dynamically adjust root growth direction and physiology, thereby optimizing resource acquisition and enhancing overall survival under heterogeneous soil conditions. Additionally, roots establish a specialized microenvironment that accommodates and selectively enriches microbial communities beneficial for root growth and protection (Uribe-Acosta et al., 2025). A key determinant of rhizosphere homeostasis is, the tightly coordinated and continuous generation, renewal, and shedding of root cap cells (Kumar & Iyer-Pascuzzi, 2020). Root-cap dynamic turnover not only facilitates root penetration through the soil but also promotes the release of border cells and exudates that shape microbial community structure and support a protective, growth-conducive environment. Conversely, impaired root cap clearance results in the retention of cap cells, leading to localized microbial accumulation (Charura et al., 2024, Rüger et al., 2024,).

The root cap secretes a substantial amount of mucilage, which serves as a nutrient source for beneficial microbes (Iijima et al., 2000). However, this interaction is selective, not all microbes can utilize this mucilage. The secreted mucilage also contains antimicrobial compounds, thereby restricting microbial colonization to those organisms capable of hydrolyzing and utilizing mucilage substrates (Hu et al., 2018, Driouich et al., 2021, Gu et al., 2016). For instance, Tran et al. (2016) demonstrated that pea root caps release DNA-based extracellular traps capable of capturing *Ralstonia solanacearum (R. solanacearum)*, although the pathogen can evade these traps through the secretion of nucleases. In addition to such physical and biochemical mechanisms, plants impose an immunity-based selection on microbial communities, thereby establishing multiple layers of control that collectively shape a beneficial microbiome (Tsai et al., 2023).

Beyond structural and biochemical defences, plant hormones play a central role in coordinating cellular communication and regulating both immunity and microbial assembly. Many pathogens exploit plant hormonal pathways by either producing phytohormones themselves or deploying effectors that manipulate host hormone signalling (Kazan, K., & Lyons, R. 2014). Cytokinin has been identified as a key regulator of root immunity, acting in conjunction with salicylic acid to mediate defence responses (Alonso-Díaz et al., 2021). Cytokinin has been reported to promote FLS2 assembly, thereby enhancing plant immune responses (Pizarro et al., 2021). Moreover, exogenous cytokinin application can increase systemic resistance in tomato (Gupta et al., 2020). In *Lotus japonicus,* cytokinin has also been shown to influence microbial colonization in roots (Soyano et al., 2024). In contrast, its antagonistic hormone auxin is often associated with increased susceptibility to infection (Kidd et al., 2011, Mutka et al., 2013). Interestingly, the root tip serves as a major regulatory hub for the integrating cytokinin and auxin signalling (Di Mambro et al., 2019).

During pathogenesis, the elongation zone and root cap are major susceptible regions, possibly due to relatively attenuated immune responses associated with their roles in beneficial microbial interactions (Tsai et al., 2023). A cell type-specific analysis of flg22 and Pep2 induced responses revealed an immune network across root tissues but did not capture root cap-specific immunity (Rich-Griffin et al., 2020). To better understand, how the root cap perceives environmental signals, Das et al. (2023) performed root cap-specific proteomics and identified spatially restricted proteins, including members of the jacalin-associated lectin (JAL) family, JAL10 and JAL20. Notably, JAL10 has been shown to mitigate stress responses under salt stress conditions. Multiple studies have demonstrated that modulation of JAL expression enhances plant resistance to pathogenic infection. For example, in rice, OsJAC1 overexpression confers broad-spectrum resistance (Weidenbach, D., et al, 2016). Similarly, elevated expression of *TaJJRLL1* has been shown to enhance resistance against fungal pathogens (Xiang, et al, 2011). However, the specific roles of Arabidopsis JAL10 and JAL20 in mediating biotic interactions have remained uncharacterized. Additionally, the molecular mechanisms underlying the role of JALs in biotic stress responses in dicots remain poorly characterized.

In this study, we demonstrate that infection by *Ralstonia pseudosolanacearum F1C1 (R. pseudosolanacearum)* induces cytokinin biosynthesis and signalling in the root tip. Elevated cytokinin levels in the root tip are associated with the downregulation of the root cap-specific genes *JAL10* and *JAL20*. Functional analyses further revealed that knockout mutants exhibit delayed pathogen invasion, whereas overexpression lines display increased susceptibility compared to Col-0. Moreover, integrated glycoproteomics and metabolomics analyses showed that cell wall biogenesis is constitutively enhanced in the mutants under basal conditions. Consistently, mutants also display increased accumulation of phenylpropanoid-derived secondary metabolites, indicating a coordinated reinforcement of structural and chemical defense mechanisms.

## Materials and Methods

### Plant material

The Arabidopsis lines *pIPT5:GFP*, *pIPT7:GFP*, *pPYK0:CKX3, ahk3/ahk4,* were provided by the Prof. Thomas Schmülling of Freie Universität Berlin. The *jal10*, *pJAL10:eGFP*, and *pJAL20:eGFP* lines were previously described by Das et al. (2023). The 35S:ARR1:GFP line was described by Zubo et al. (2017). For the generation of JAL10 and JAL20 overexpression lines, coding sequences were amplified with attB overhangs, cloned into pDONR221 using BP recombination, and transferred into pK2GW7. Constructs were transformed into *Agrobacterium tumefaciens GV3101* and introduced into Arabidopsis by floral dip. Independent transgenic lines were selected on kanamycin, and two lines with comparable expression were used for further analyses. Knockout mutants were obtained from NASC and confirmed by genotyping.

### Plant growth conditions

Arabidopsis seeds were surface-sterilized with 70% ethanol followed by 4% sodium hypochlorite and washed three times with sterile water. Seeds were plated on half-strength MS medium containing 1% sucrose and 1% agar (pH 5.7), stratified at 4°C in darkness for 72 h, and grown for 5 days under long-day conditions (16 h light/8 h dark, 150 μmol m⁻² s⁻¹, 22°C/20°C day/night, 65% relative humidity).

### Infection assays

*Ralstonia pseudosolanacearum* F1C1 was cultured in BG broth as previously described (Kumar et al., 2017). Bacterial cultures were collected by centrifugation, washed twice with sterile water, and resuspended to the required OD₆₀₀. For promoter activity assays, 5-day-old seedlings were transferred to half-strength MS medium lacking sucrose and MES. *R. pseudosolanacearum-mCherry* was adjusted to OD₆₀₀ = 0.01, and 5 μL of inoculum was applied directly to the root tip. Plates were incubated at 28°C, and fluorescence was monitored at the indicated time points using Leica SP8 and Olympus FV3000 confocal microscopes. The detailed protocols for the invasion assay, CFU assay, and soil-drench assay were provided in the Supplemental Information (SI).

### Chip-qPCR

Chromatin immunoprecipitation (ChIP) assays were performed using approximately 2 g of root tissue from 10-day-old *Arabidopsis thaliana* seedlings expressing *35S:ARR1:GFP* (Col-0) and *35S:eGFP* (Col-0), grown on half-strength MS medium. Prior to crosslinking, seedlings were treated with 1 µM 6-benzylaminopurine (BAP) for 1 hour to activate cytokinin signalling. Crosslinking was carried out using 1% formaldehyde to stabilize protein–DNA interactions, and the reaction was subsequently quenched. Chromatin was then isolated and fragmented by sonication using a Qsonica 800R sonicator, by applying 30 cycles of 15-second pulses at 70% amplitude, with 45-second intervals between pulses, to obtain appropriately sheared DNA fragments. ARR1-DNA complexes were immunoprecipitated using protein G magnetic beads (Dynabeads, 10004D) conjugated with an anti-GFP antibody (Abcam, ab290), while rabbit IgG was used as a negative control to assess non-specific binding. Following immunoprecipitation, the beads were extensively washed to remove non-specifically bound material, and the protein–DNA complexes were eluted. Crosslinks were then reversed to release the associated DNA. The enriched DNA was subsequently analysed by quantitative PCR (qPCR) using primer sets listed in Table 1. Enrichment levels were calculated relative to the IgG control, allowing quantification of ARR1 binding at the target promoter regions.

### Glycoproteomics and Metabolomics

Root tip glycoproteins were extracted from infected and mock-treated seedlings, digested with trypsin, and analysed using Waters LC/MS G2-XS Q-TOF (Milford, Massachusetts, USA). Differentially abundant proteins were identified by quantitative glycoproteomic analysis followed by Gene Ontology enrichment. Similarly, metabolites were extracted from root tips collected at 48 hpi and analysed using LC–MS/MS. Data processing, metabolite annotation, and statistical analyses were performed using MS-DIAL and MetaboAnalyst. The detailed protocol for Glycoproteomics and Metabolomics is provided in the Supplemental Information (SI).

### VIGS-mediated silencing and phenotypic analysis in *Solanum lycopersicum*

The TRV based vectors pTRV1 and pTRV0 were obtained from Chandan, R. K., et 2023, NIPGR, New Delhi. The target region for VIGS silencing of *SLJAL1, SLJAL5* and *SLJAL9* approximately 300 base pairs in length was predicted by SGN VIGS (http://vigs.solgenomics.net) The predicted regions were PCR amplified using specific target primer pairs, and cloned into multiple cloning sites of the pTRV2 vector and confirmed by Sanger sequencing. The constructs were then transformed into *Agrobacterium tumefaciens* GV3101 cells. Subsequently the cultures of pTRV:*SlJAL1,* pTRV:*SlJAL5* and pTRV:*SlJAL9* were mixed with pTRV1 at 1:1 ratio(OD_600_∼0.1) and infiltrated into 2-week-old tomato seedlings. SlPDS is used as a positive control for knock-down. The silencing of *pTRV:SlJAL1, pTRV:SlJAL5* and *pTRV*:*SlJAL9* was confirmed using RT-qPCR. For analysing the phenotype of VIGS-silenced plants, *R. pseudosolanacearum* of OD_600_=0.1 was inoculated into the pots of 3-week-old tomato plants using the drench method. The plants were maintained in a greenhouse at 28°C under a 16:8 h light/dark cycle with 80% humidity. Disease symptoms were observed for seven days after inoculation and photographed using a DSLR camera. Each experiment included at least 15 infected plants and was independently replicated. The wilting index was determined following the method described by Hiles et al., 2024. Briefly, wilting symptoms were assessed using a scale from 0 to 4, where 0 indicated no visible symptoms,1 indicated ≤25% wilting symptoms, 2 indicated ≤50% wilting symptoms, 3 indicated ≤75% wilting symptoms and 4 indicated ≤100% wilting of leaves. Based on these wilt scores, the disease index (DI) was calculated using the formula: DI = Nw/N (Nw = number of leaves wilted, N = total number of leaves). For CFU quantification, wilted samples were tested on the 7^th^ day as mentioned above.

### RT-qPCR

To quantify transcript levels in *Arabidopsis thaliana* and *Solanum lycopersicum*, total RNA was extracted from three independent biological replicates of both infected and control samples. RNA isolation was performed using the Takara RNA extraction kit (Cat. No. 6110A) according to the manufacturer’s instructions. For cDNA synthesis, 1000 ng of total RNA was used for reverse transcription. The resulting cDNA was subsequently used for quantitative real-time PCR (RT–qPCR) to assess transcript abundance, following the protocol described by Ramireddy et al. (2018). The analysis was conducted using three biological replicates to ensure reproducibility.

## Results

### Cytokinin biosynthesis is induced in the root tip, and its signalling is required for root immunity

Cytokinin and auxin signalling play important roles in regulating root responses during infection by *R. pseudosolanacearum*. Previous studies have implicated cytokinin in root immunity; however, its spatial regulation and functional contribution in the root tip remain incompletely understood. (Alonso-Díaz et al., 2021, Pizarro, et al., 2021, Zhang et al., 2024, Wang et al., 2024). To address this, we examined cytokinin dynamics in the root tip, a primary site of cytokinin biosynthesis following infection with *R. pseudosolanacearum*. Cytokinin levels were monitored using the reporter lines *pIPT5:GFP* and *pIPT7:GFP*, which mark sites of cytokinin biosynthesis in the root cap and meristematic zone (Miyawaki et al 2004; Figure 1A-B&D). After infection, both IPT5 and IPT7 activities significantly increased at 24 and 48 hours, indicating enhanced cytokinin production in the root tip. Simultaneously, auxin levels decreased markedly at 48 hours, as indicated by increased expression of the DII-VENUS reporter, suggesting a hormonal shift favouring cytokinin (Figure 1C&D).

**Figure 1.**
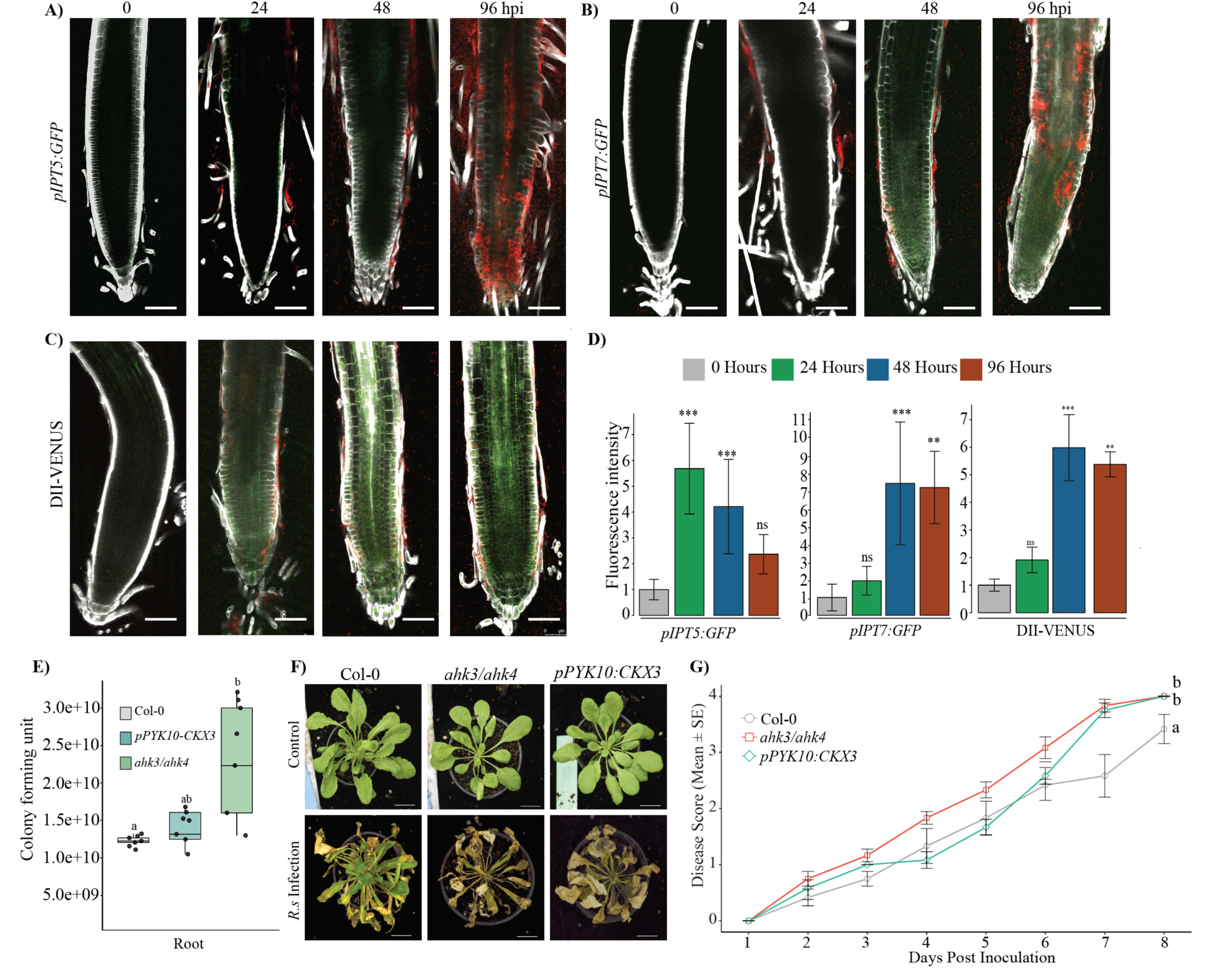
Cytokinin biosynthesis is induced during root infection and signaling is required for root immunity. (A, B) Confocal images showing increased promoter activity of (A) pIPT5 and (B) pIPT7 following infection with *R. pseudosolanacearum*. GFP fluorescence indicates cytokinin biosynthetic activity. (C) Auxin distribution monitored using the DII-VENUS reporter, in which fluorescence intensity is inversely proportional to auxin levels. (D) Quantification of fluorescence intensity in infected roots. Statistical significance was determined using the Kruskal–Wallis test followed by Dunn’s multiple comparison test. Asterisks indicate significant differences (*P ≤ 0.05, **P ≤ 0.01, ***P ≤ 0.001). Pathogen fluorescence is shown in red, GFP in green, and cell walls were stained with calcofluor white. Scale bars, 100 μm. (E) Quantification of bacterial colony-forming units (CFUs) following inoculation with *R. pseudosolanacearum* (OD_600_ = 0.01). Statistical significance was determined using Student’s *t*-test. (F, G) Disease progression in cytokinin-related mutants following soil-drench inoculation. (F) Representative phenotypes and (G) wilting index over time. Statistical analyses were performed using Friedman and Duncan tests for disease progression and Wilcoxon signed-rank tests for genotype comparisons. Different letters indicate significant differences at 8 dpi. Detailed statistical analysis with replication and sample size are provided in Table S1. Scale bar, 1 cm.

To assess the functional contribution of cytokinin signalling to root immunity, we tested the cytokinin receptor mutant *ahk3/ahk4* (Riefler et al., 2006), and the *pPYK10-CKX3* (Werner, T., et al 2010), which has reduced cytokinin levels specifically in roots, against *R. pseudosolanacearum*. Colony-forming unit (CFU) assays revealed significantly higher bacterial accumulation in *ahk3/ahk4* mutants, followed by *pPYK10:CKX3*, compared with Col-0 three days post-inoculation (Figure 1E). Consistently, soil-drench infection assays showed that *ahk3/ahk4* plants exhibited the most severe disease symptoms, whereas Col-0 displayed comparatively reduced susceptibility (Figure 1F&G). These findings indicate that pathogen infection boosts cytokinin production in the root tip and that effective cytokinin perception is crucial for root immunity. The greater susceptibility in cytokinin perception mutants emphasizes the importance of signalling in defence responses. Given that the root cap serves as a major interface for microbial interactions and a key site of cytokinin regulation, we sought to identify the molecular players that mediate root cap immunity.

### Cytokinin directly represses JAL10 and JAL20 via type-B transcription factor ARR1

To identify root cap-specific factors regulated during infection, we focused on jacalin-associated lectins JAL10 and JAL20, previously identified as spatially restricted proteins in the root cap (Das et al., 2023). To determine their transcriptional response to *R. pseudosolanacearum* infection, we analysed promoter activity using *pJAL10:eGFP* and *pJAL20:eGF*P reporter lines in Arabidopsis seedlings. Under control conditions, *pJAL10:eGFP* expression is specifically observed in the root cap, including both the columella and lateral root cap. *pJAL20:eGFP* is predominantly expressed in the lateral root cap, extending into the meristematic zone, reaching till the elongation zone and shows strong expression in the root-root cap of the lateral root as well (Figure 2). However, upon infection with *R. pseudosolanacearum*, the intensity of fluorescence from both reporters decreased progressively over 96 hours, suggesting that the transcription of these genes was being repressed (Figure 2 A-C). Notably, this reduction initiated in the lateral root cap and gradually extended toward the columella, suggesting spatially coordinated regulation during infection. Consistently, a similar decline in reporter activity was also observed in *pJAL8:eGFP,* the closest homolog of *JAL10* and *JAL20,* upon infection with bacteria, (Figure S1A).

**Figure 2.**
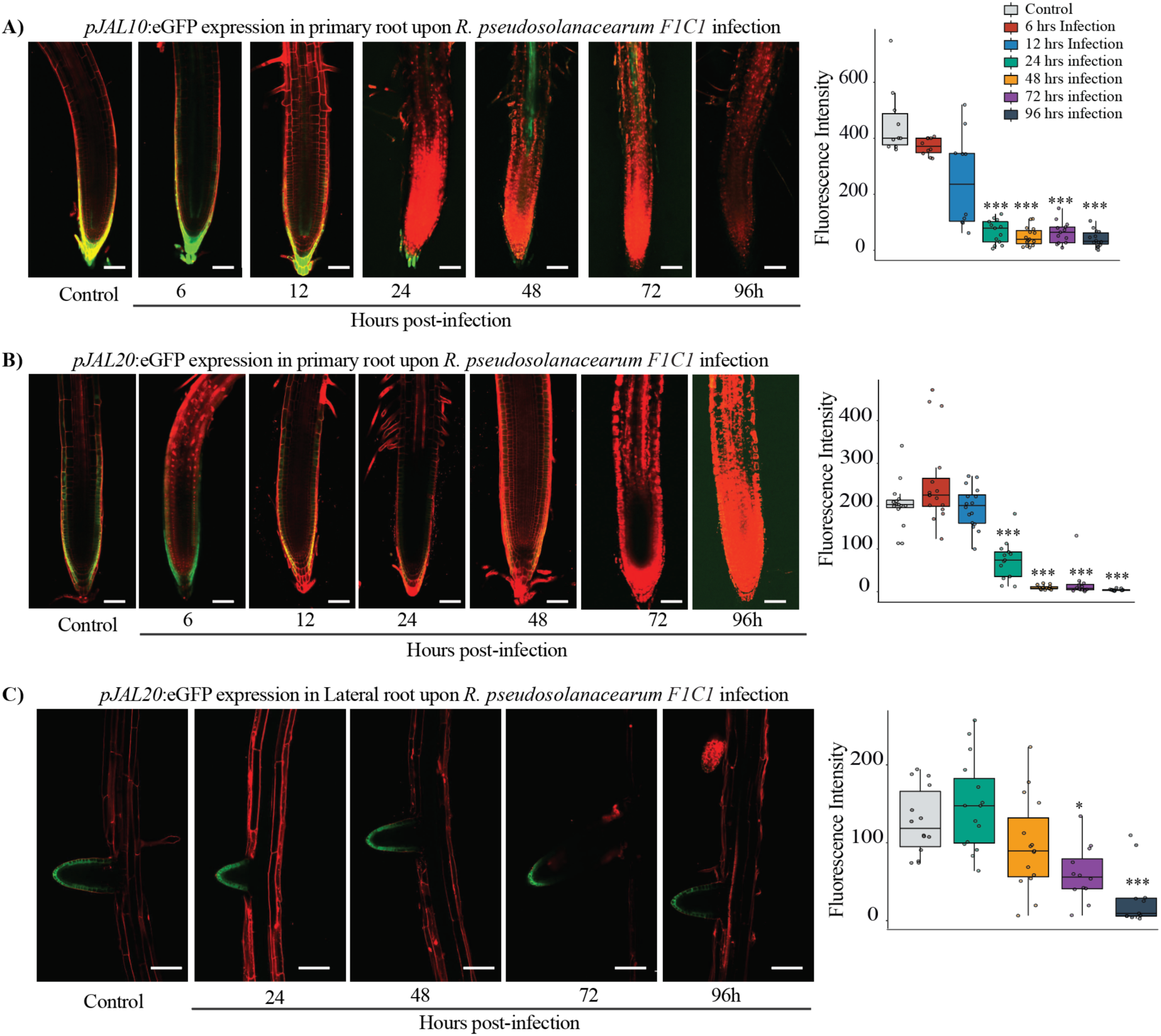
Transcript levels of *JAL10* and *JAL20* are repressed during *R. pseudosolanacearum* infection. (A-C) Confocal images showing reporter activity of (A) *pJAL10*, (B) *pJAL20* in the primary root cap, and (C) *pJAL20* in root cap of lateral root during infection with *R. pseudosolanacearum* over a 96-hour time course. Mock-treated plants served as controls. GFP fluorescence is shown in green and cell walls stained with propidium iodide in red. Scale bars, 100 μm. Fluorescence intensity was quantified from three independent biological replicates (n = 15 per experiment). Statistical significance was determined using the Kruskal–Wallis test followed by Dunn’s multiple comparison test (*P ≤ 0.05, **P ≤ 0.01, ***P ≤ 0.001).

To determine whether cytokinin signalling was involved in this repression, we analysed the promoter regions of *JAL10* and *JAL20*. We identified several type-B ARR-binding motifs, suggesting potential regulation by cytokinin transcription factors. When we applied 6-benzylaminopurine (BAP), a synthetic cytokinin, exogenously for 30, 60, and 120 minutes, we observed rapid suppression of reporter activity within 30 minutes for JAL10 and 60 minutes for JAL20 (Figure 3A-C). Interestingly, a similar analysis with exogenous application of synthetic auxin (IBA) reveals that *JALs* are positively regulated by auxin treatment (Figure S2A-B). Further the transcript levels of *JAL10* and *JAL20* in the auxin receptor mutant *tir1* is reduced under control conditions suggest that auxin might play a positive regulator of JALs (Figure S2C). These results suggest that cytokinin signalling can rapidly modulate the transcriptional activity of *JAL10* and *JAL20* in the root cap, potentially fine-tuning their response during pathogen infection.

**Figure 3.**
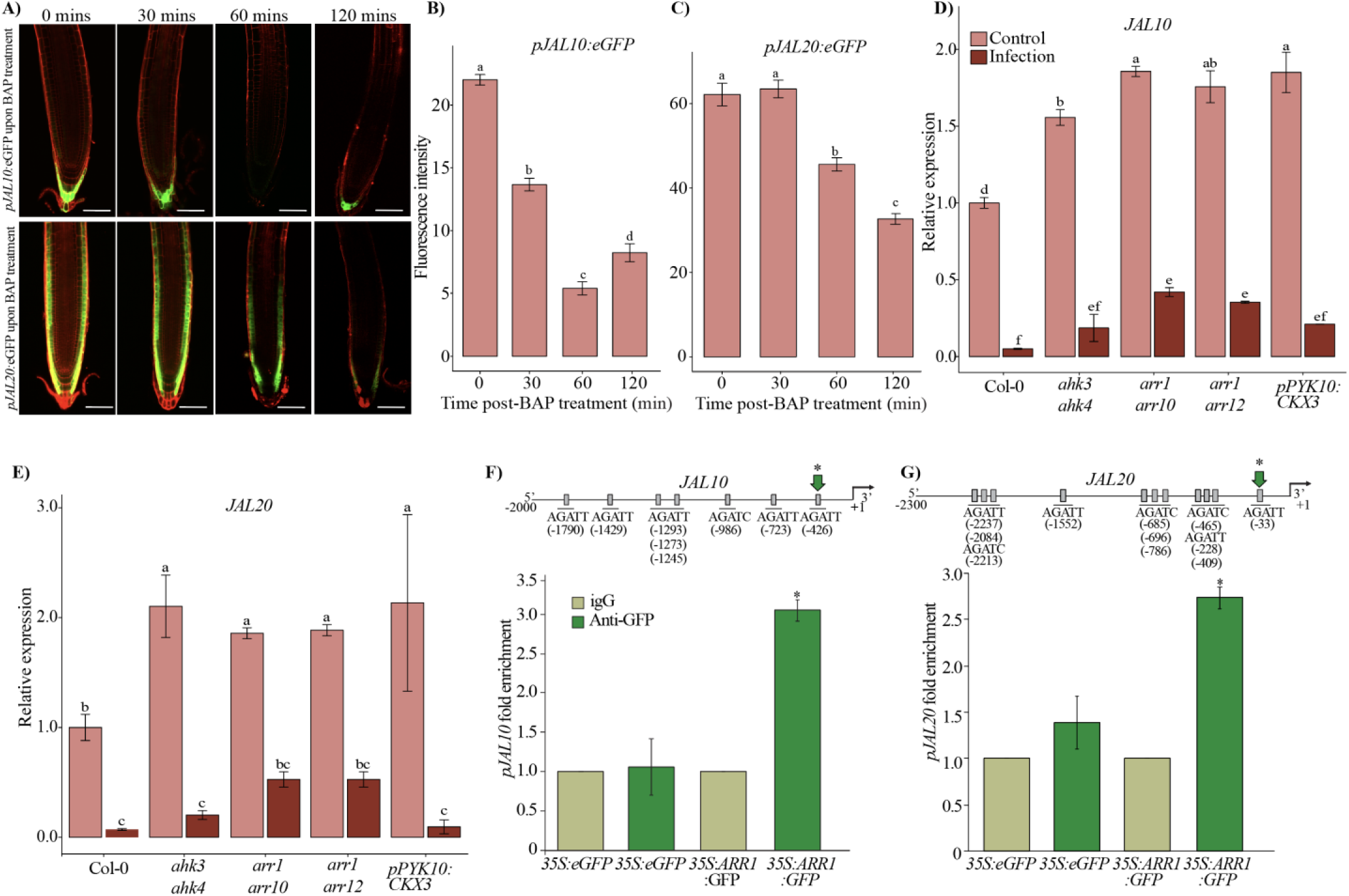
Cytokinin negatively regulates the expression of JAL10 and JAL20. (A) Confocal images showing *pJAL10* and *pJAL20* reporter activity following treatment with 1 μM BAP. GFP fluorescence is shown in green and cell walls stained with propidium iodide in red. (B, C) Quantification of fluorescence intensity following cytokinin treatment. Data represent three independent biological replicates (n = 15 per experiment). (D, E) Relative transcript levels of *JAL10* and *JAL2*0 in cytokinin receptor mutants (*ahk3/ahk4*), signaling mutants (*arr1/arr10* and *arr1/arr12*), and root specific cytokinin-deficient lines (*pPYK10:CKX3*) under control and infection conditions. (F, G) ChIP–qPCR analysis showing ARR1 enrichment at promoter regions of JAL10 and JAL20. Green arrows indicate regions with significant ARR1 binding. Numbers indicate distance upstream of the transcription start site. Error bars represent ± SEM. Statistical significance was determined using two-way ANOVA followed by Fisher’s LSD test or Student’s *t*-test where appropriate (*P ≤ 0.05).

Next, we investigated JAL10 and JAL20 expression across various genetic backgrounds associated with cytokinin signalling. In cytokinin receptors (*ahk3/ahk4*), signalling (*arr1/arr10* and *arr1/arr12*) mutants, and cytokinin-deficient (*pPYK10:CKX3*) lines, transcript levels of *JAL10* and *JAL20* were elevated under basal conditions compared to Col-0. Upon *R. pseudosolanacearum* infection, transcript levels were reduced across all genotypes, reflecting the pathogenesis-induced repression of these genes. However, the signalling mutants *arr1/arr10* and *arr1/arr12* retained higher transcript abundance relative to the other mutants (Figure 3D-E). This suggests that the type-B ARRs are required for efficient downregulation of *JAL10* and *JAL20*. To test whether type-B ARRs directly regulate JAL transcription, we performed chromatin immunoprecipitation (ChIP) followed by qPCR using ARR1:GFP. Strong enrichment of ARR1 binding was detected at promoter regions proximal to the transcription start sites of both *JAL10* (at −426 bp) and J*AL20* (at −33 bp) (Figure 3F-G). These findings confirm that cytokinin signalling directly represses transcription of *JAL* genes by binding ARRs to their promoters. Together, these results establish JAL10 and JAL20 as direct targets of cytokinin signalling and reveal a mechanism by which pathogen-induced hormonal changes modulate root cap gene expression.

### JAL10 and JAL20 negatively regulate root immunity

To better understand the functional implications of JAL10 and JAL20 on root immunity, we investigated how *loss-of-function* mutants and overexpression lines responded during infections with *R. pseudosolanacearum*. We generated overexpression lines under the 35S promoter and confirmed the knockout mutants through genotyping and transcript analysis (Figure S3A). We performed *in vitro* invasion assays using mCherry-labelled *R. pseudosolanacearum* to track pathogen invasion of roots at 24, 48, 72, 96 and 120 hours (Figure S3B-C). The spatial progression of bacterial penetration was monitored at the root tip and elongation zone at 24-hour intervals up to 120 hpi (Figure S3C), as *R. pseudosolanacearum* predominantly accumulates in these regions (Zhang et al., 2024). In wild-type plants, bacterial accumulation was first detected at the root tip and elongation zone at 24 hours post-inoculation and increased progressively over time. By 48 hpi, strong fluorescence signals were observed surrounding these regions, indicating substantial bacterial accumulation in the root tip and elongation zone (Figure S3C). In the JAL overexpression lines, we observed a notable increase in bacterial accumulation, with strong fluorescence not just at the root tip and elongation zone but also spreading into vascular tissues, indicating rapid invasion and systemic spread of the pathogen (Figure 4A-H). Conversely, the *jal10* and *jal20* mutants displayed delayed pathogen spread, with reduced bacterial entry and slower colonisation throughout the observed period. Quantitative analyses of the fluorescence intensity at 120 hpi corroborated these observations. The *jal10* and *jal20* mutants had significantly lower bacterial loads in the root tip than Col-0, while the overexpression lines showed greater colonisation. Similar trends were observed in the elongation zone and vascular tissues where the mutants consistently had lower pathogen loads, whereas the overexpression lines showed heightened susceptibility. Notably, in the vasculature, a critical niche for the systemic spread of *R. pseudosolanacearum* fluorescence in the mutants was reduced by ∼30% in *jal10* and ∼40% in *jal20* relative to Col-0. In contrast, JAL10 overexpression lines exhibited a 70-100% increase, while JAL20 overexpression lines showed a 10-20% increase (Figure 4A-H). Furthermore, we conducted colony-forming unit (CFU) assays to quantify bacterial colonisation. Consistent with our imaging results, the *jal10* mutants exhibited reduced bacterial loads in both roots and shoots compared to Col-0, and the overexpression lines showed increased colonisation levels. A similar pattern emerged for JAL20, with mutants showing lower bacterial accumulation and overexpression lines revealing elevated levels (Figure 4I-J, Figure S4A).

**Figure 4.**
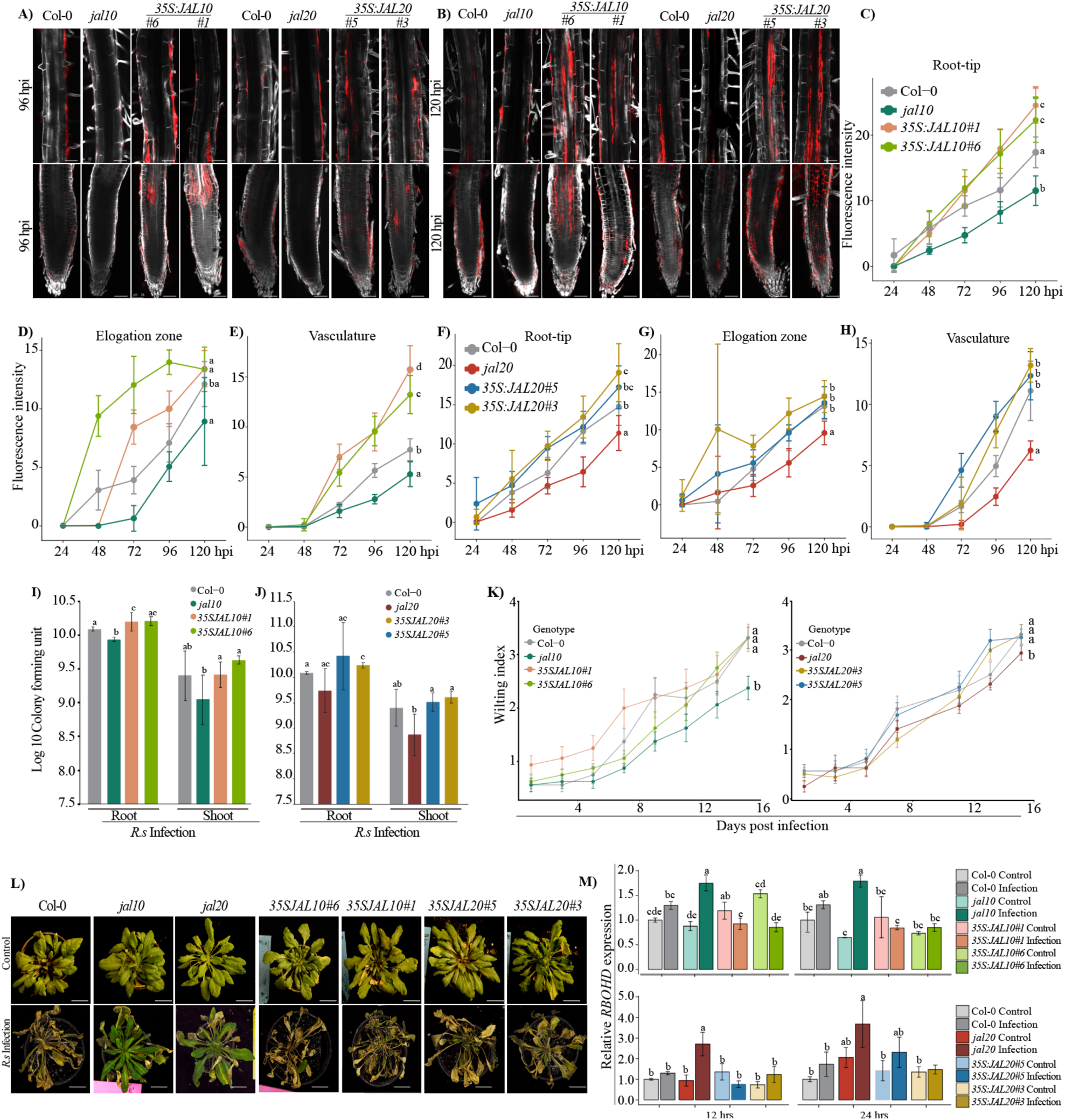
JAL10 and JAL20 negatively regulate root immunity. (A, B) Confocal images showing progression of *R. pseudosolanacearum-mCherry* during root invasion assays at 96 and 120 hpi in JAL10 and JAL20 functional mutants. Bacterial fluorescence is shown in red and cell walls stained with calcofluor white. Scale bars, 50 μm. (C–H) Quantification of bacterial fluorescence intensity in the root cap, elongation zone, and vasculature of JAL10 and JAL20 functional mutants. Statistical analyses were performed using Friedman tests followed by Dunn’s multiple comparison tests for temporal progression and Wilcoxon tests for genotype comparisons. Different letters indicate significant differences at 120 hpi. Corresponding Statistical details are provided in Tables S2 and S3. (I, J) Quantification of bacterial colony-forming units (CFUs) in roots and shoots of JAL10 and JAL20 functional mutants following infection. (K, L) Disease progression following soil-drench inoculation. (K) Wilting index over time and (L) representative disease phenotypes. Statistical analyses were performed using Kruskal–Wallis and Wilcoxon rank-sum tests. Corresponding statistical details are provided in Tables S4 and S5. Scale bar, 1 cm. (M) Relative transcript levels of *RBOHD* in JAL10 and JAL20 functional mutants under control and infection conditions. Statistical significance was determined using two-way ANOVA followed by Fisher’s LSD test.

Lastly, we evaluated disease progression through soil-drench infection assays. While all genotypes ultimately developed symptoms, *jal10* and *jal20* mutants experienced significantly delayed disease progression compared to both Col-0 and overexpression lines. In contrast, the overexpression lines exhibited accelerated wilting and signs of heightened disease severity (Figure 4K-L). Consistently with the role of reactive oxygen species (ROS) in plant defence, infection by *R. solanacearum* is also associated with a pronounced increase in ROS accumulation (Flores-Cruz, Z., & Allen, C. 2009). We analysed expression of the NADPH oxidase gene *RBOHD* following infection. Transcript levels of RBOHD were significantly elevated in *jal10* and *jal20* mutants at early time points, indicating enhanced activation of ROS-mediated defence responses (Figure 4M). Together, the results obtained from the three independent infection methods, *in vitro* invasion assay, quantification of CFUs and the soil-drenching infection assay demonstrate that JAL10 and JAL20 act as negative regulators of root immunity. Their repression during infection enhances defence responses, whereas elevated expression promotes pathogen colonisation and disease progression.

### Loss-of-JAL function leads to increased accumulation of cell wall-related glycoproteins

To investigate the molecular basis underlying the altered infection phenotypes of JAL mutants, we performed glycoproteomic profiling of wild-type plants, knockout mutants, and overexpression lines under control and infection conditions. Because JAL10 and JAL20 localize to the endoplasmic reticulum (Das et al., 2023), we hypothesized that they may influence protein processing or glycosylation. Glycoproteomic analysis revealed extensive protein modifications (Tables S6–S8), including 257 O-glycosylation sites, 186 N-glycosylation sites, and 30 C-mannosylation events, together with carbamidomethylated cysteines and oxidized methionines (Tables S8). In total, 501 modified proteins were identified, highlighting the diversity of peptide modification states. Functional enrichment analysis revealed strong enrichment of proteins associated with cell wall organization, glycan modification, and secondary metabolism (Figure 5A-C). Based on the selection criteria described in the glycoproteomic analysis section, 52 and 191 significantly altered proteins were identified in JAL10 and JAL20 functional mutants, respectively, relative to Col-0 (Figure S6A-B).

**Figure 5.**
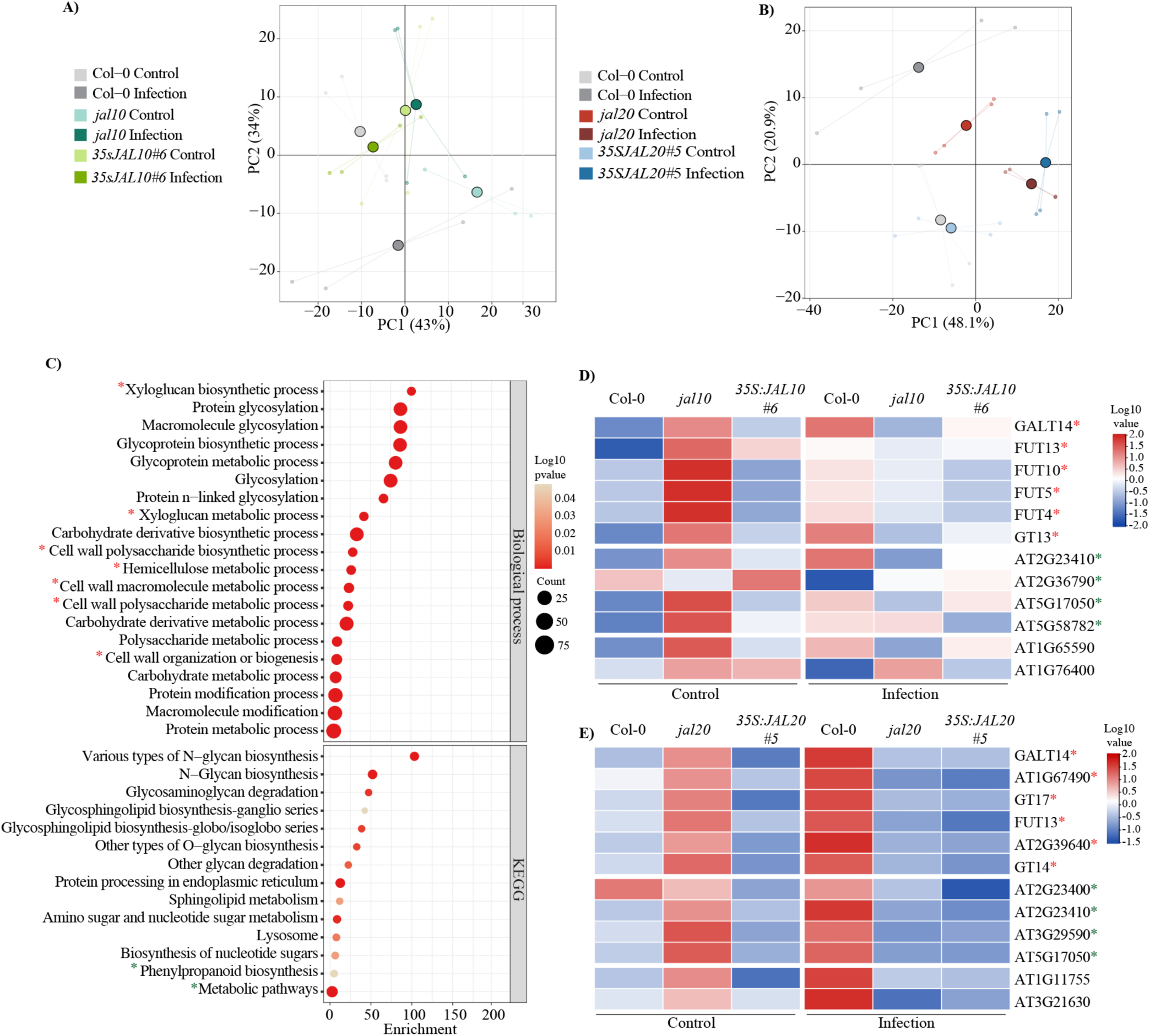
Loss of JAL function enhances accumulation of cell wall-associated glycoproteins. (A, B) Principal component analysis of glycoproteomic profiles from JAL10 and JAL20 functional mutants. (C) Gene Ontology enrichment analysis showing overrepresentation of cell wall-associated biological processes in the glycoproteome. (D, E) Heatmaps showing relative abundance of proteins associated with cell wall biogenesis, phenylpropanoid metabolism, and stress responses. Red asterisks indicate cell wall-associated proteins, and green asterisks indicate proteins involved in secondary metabolism.

Under control conditions, *jal10* mutants exhibited the highest abundance of proteins that were significantly altered when compared to Col-0 and JAL10 overexpression genotypes (Figure S6C-D). In contrast, protein abundance in JAL20 functional lines was more evenly distributed across genotypes, with a slight increase in the overexpression line (Figure S6E-F). In both mutant backgrounds, enriched proteins were predominantly associated with glycosylation-related processes, consistent with the enrichment strategy, and were more strongly associated with cell wall-related functions than in Col-0 and overexpression lines (Figure S6G and I). Upon *R. pseudosolanacearum* infection, the abundance of significantly altered proteins shifted predominantly toward Col-0, accompanied by increased accumulation of proteins involved in cell wall biogenesis. In contrast, JAL10 and JAL20 overexpression lines showed reduced accumulation of proteins associated with defence responses and secondary metabolism during infection (Figure S6H and J). These findings suggest that elevated JAL expression suppresses glycoprotein accumulation associated with cell wall remodelling and stress adaptation.

*k*-means clustering analysis further supported these patterns (Figures S8-S9). In JAL10 functional mutants, clusters 2, 3, and 4 showed strong enrichment in the knockout line under basal conditions and resembled the infection-responsive profile of infected Col-0 plants. These clusters included proteins associated with cell wall biogenesis and glycan modification, including galactosyltransferases (GALT2, GALT5, and GALT9) and fucosyltransferases (FUT4, FUT5, FUT10, FUT11, and FUT13), which are involved in cell wall assembly and defence responses (Sarria et al., 2001; Strasser et al., 2007; Zhang et al., 2019). Proteins associated with secondary metabolism, including UGT78D2 and cis-prenyltransferases involved in flavonoid and isoprenoid biosynthesis, were also enriched. Several stress-responsive proteins clustered within these groups, indicating that loss of JAL10 affects multiple pathways involved in cell wall remodelling and stress adaptation. A similar trend was observed in JAL20 functional mutants, where the basal expression profile of *jal20* resembled that of infected Col-0 plants. Cluster 4 showed strong enrichment of galactosyltransferase and fucosyltransferase family proteins, suggesting constitutive activation of cell wall biosynthesis and remodelling pathways.

Collectively, these glycoproteomic data indicate that loss of JAL10 and JAL20 function promotes a preconditioned defence state characterized by enhanced accumulation of cell wall-associated glycoproteins, stress-responsive proteins, and secondary metabolism-related proteins. This constitutive reinforcement of structural and biochemical defences likely contributes to enhanced basal resistance and delayed pathogen invasion.

### JAL mutants show increased phenylpropanoid metabolism and defence-related compounds

To complement the glycoproteomic analysis and assess downstream metabolic changes, we performed untargeted metabolomic profiling of JAL functional mutants and wild-type plants under infection conditions. Our downstream analysis identified 4,811 and 3,399 significant metabolic features in the *JAL10* and *JAL20* functional mutant backgrounds (including comparisons with Col-0), respectively (Figure S10A-B). Among these, 313 metabolites in *JAL10* and 374 metabolites in *JAL20* were successfully annotated as significantly altered compounds. These annotated metabolites were subjected to partial least squares discriminant analysis (PLS-DA) to prioritize features contributing to genotype-specific metabolic variation (Supplemental figure 10 C-D). Our analyses showed that *jal* mutants were particularly enriched in metabolites from the phenylpropanoid pathway, which is crucial for plant defence and generates compounds that bolster antimicrobial activity and support cell wall integrity.

Phenylalanine is a critical aromatic amino acid and a central precursor of the phenylpropanoid pathway (Figure 6A). Following infection, we observed a significant increase in key upstream intermediates, such as phenylalanine and trans-cinnamic acid, in the *jal* mutants, indicated a heightened activation of the phenylpropanoid pathway (Figure 6B-C). These metabolites play important roles in plant defence by contributing to antimicrobial compound production and reinforcing cell wall structures (Singh, 2025). Further, we also saw increased levels of downstream metabolites, such as flavonoids like kaempferol and quercetin, and a coumarin derivative, sideretin, and leucodelphinidin, an important intermediate required for anthocyanin biosynthesis were enriched in the *jal10* and *jal20* compared to Col-0 and JAL overexpressing lines indicating more biosynthesis of antioxidant, ROS scavenging, defence-related compounds and helps in cell wall strengthening (Sia, et al., 2013; Liu, et al., 2021). Moreover, we found that *jal* mutants accumulated higher levels of lignin-related metabolites, such as sinapate derivatives and syringin. These compounds are precursors for lignin synthesis, which enhances cell wall rigidity and helps resist pathogen attacks (König, S., et al, 2014, Pascual, M. B., et al, 2016). This aligns well with the idea that *jal* mutants have strengthened structural defences. In addition to the phenylpropanoid metabolites, the *jal* mutants had more oxylipin derivatives, such as 9,10-epoxyoctadecatrienoic acid, which play a role in callose deposition and cell wall fortification. We also observed elevated levels of indole-derived metabolites linked to glucosinolate production, suggesting the activation of additional chemical defence pathways. Overall, these results suggest that *loss-of-function* of *jal10 and jal20* in plants, undergoes significant metabolic reprogramming that bolsters their chemical and structural defences against pathogens.

**Figure 6.**
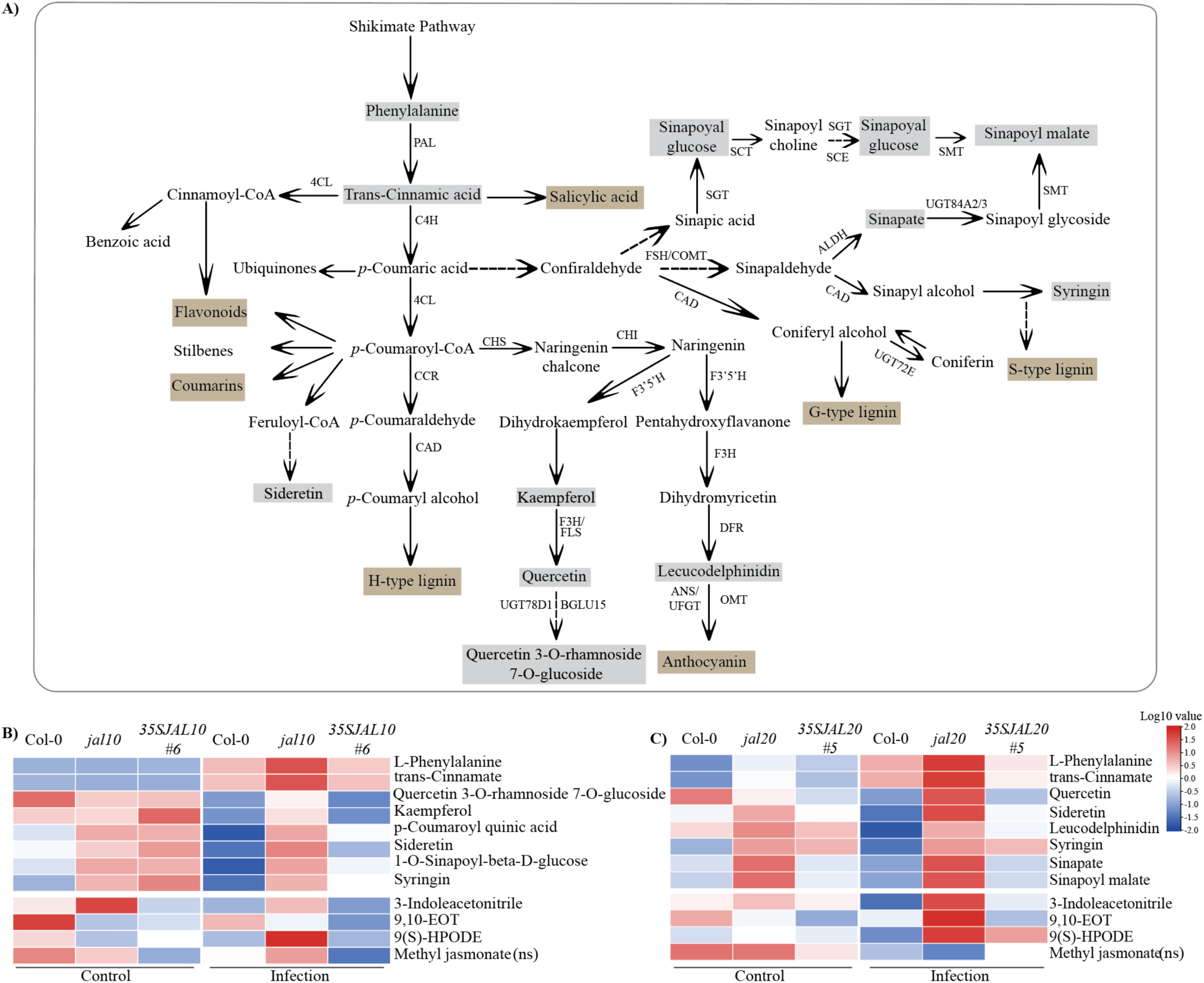
JAL mutants exhibit enhanced accumulation of phenylpropanoid-derived defense metabolites. (A) Schematic overview of the phenylpropanoid pathway and associated defense branches. Grey boxes indicate metabolites enriched in *jal10* and *jal20* mutants following infection with *R. pseudosolanacearum*, whereas beige boxes indicate terminal defense-related products. (B, C) Heatmaps showing significantly enriched phenylpropanoid-associated metabolites in JAL10 and JAL20 functional mutants at 48 hpi. Metabolites associated with lignin biosynthesis, flavonoid metabolism, and defense responses are prominently enriched in knockout lines. Phenylalanine ammonia-lyase (PAL), Cinnamate 4-hydroxylase (C4H), 4-coumarate:CoA ligase (4CL), Cinnamoyl-CoA reductase (CCR), Cinnamyl-alcohol dehydrogenase (CAD), Chalcone synthase (CHS), Chalcone isomerase (CHI), Flavanone 3′,5′-hydroxylase (F3′5′H), Flavanone 3-hydroxylase (F3H), Flavonol synthase (FLS), Dihydroflavonol 4-reductase (DFR), Anthocyanidin synthase (ANS), UDP-glucose flavonoid 3-O-glucosyltransferase (UFGT), O-methyl transferases (OMT), UDP-glycosyl transferase (UGT), Caffeic acid 3-Omethyltransferase (COMT), Aldehyde dehydrogenase (ALDH), UDP-glucose:sinapate glucosyltransferase (SGT), Sinapoylglucose:choline sinapoyltransferase (SCT), Sinapine esterase (SCE), Sinapoylglucose:malate sinapoyltransferase (SMT).

### Silencing of Solanum lycopersicum JAL orthologs enhances tolerance to R. pseudosolanacearum

In our quest to determine whether the regulatory function of JAL proteins is conserved across species, we focused on identifying JAL orthologs in tomato (*Solanum lycopersicum*). Through our domain-based structure analysis, we identified nine high-confidence tomato proteins containing the JAL domain (Figure S11A). For further characterisation, three candidate genes were selected, *Solyc01g00620 (SLJAL1)*, *Solyc09g083020 (SLJAL5)*, and *Solyc10g078600 (SLJAL9)* based on their highest expression in the root as per the transcriptome data. Validation of tissue-specific expression by RT-qPCR revealed that these JAL genes are expressed throughout the plant, with particularly high expression detected in the roots (Figure S11B-D). To explore how these proteins are involved in the plant’s response to infection by *R. pseudosolanacearum*, we examined different tomato varieties that have varying levels of tolerance to this pathogen (Acharya et al., 2018; Kumar S, et al., 2018). Interestingly, we found that in the highly tolerant variety (Utkala Kumari) and moderate susceptible variety (Arka Vikas), JAL gene expression decreased after infection. In contrast, the highly susceptible variety Pusa Ruby showed a strong increase in JAL transcripts (Figure S11E). This inverse relationship between JAL expression and disease tolerance suggests that these proteins may play a significant role in determining the plant’s susceptibility to bacterial wilt disease. To directly test this, we employed virus-induced gene silencing (VIGS) to suppress expression of selected tomato JALs in the susceptible Pusa Ruby genotype background. Phytoene Desaturase (PDS) was included as a positive control, and approximately 20 days after VIGS infiltration, the characteristic photobleaching phenotype was observed in pTRV:PDS plants, confirming successful VIGS infection (Figure S11F). Furthermore, transcript quantification in VIGS plants showed that the selected genes were significantly downregulated, indicating effective silencing of the target transcripts. (Figure S11G). After confirming the downregulation of the target genes, a soil-drench infection assay was performed using *R. pseudosolanacearum* (Figure 7A). Disease progression was monitored for seven days after infection by recording the wilting index. Plants carrying the pTRV:0 vector control showed clear wilting symptoms, whereas plants with SlJAL1 and SlJAL5 silenced showed reduced wilting. SlJAL9-silenced plants showed the lowest disease symptoms (Figure 7B). We further quantified the CFU. The bacterial population in the roots did not differ significantly between the lines. However, in the shoots, fewer colonies were observed only in SlJAL9-silenced plants (Figure 7C). In addition, the defence-related genes *PR1a, PR1b, SD2*, and *WRKY75* showed higher expression after *R. pseudosolanacearum* Infection. Among the silenced lines, SlJAL5 showed the strongest induction of these defence genes, while *PR1b* expression was particularly higher in SlJAL9-silenced plants (Figure 7D-G).

**Figure 7.**
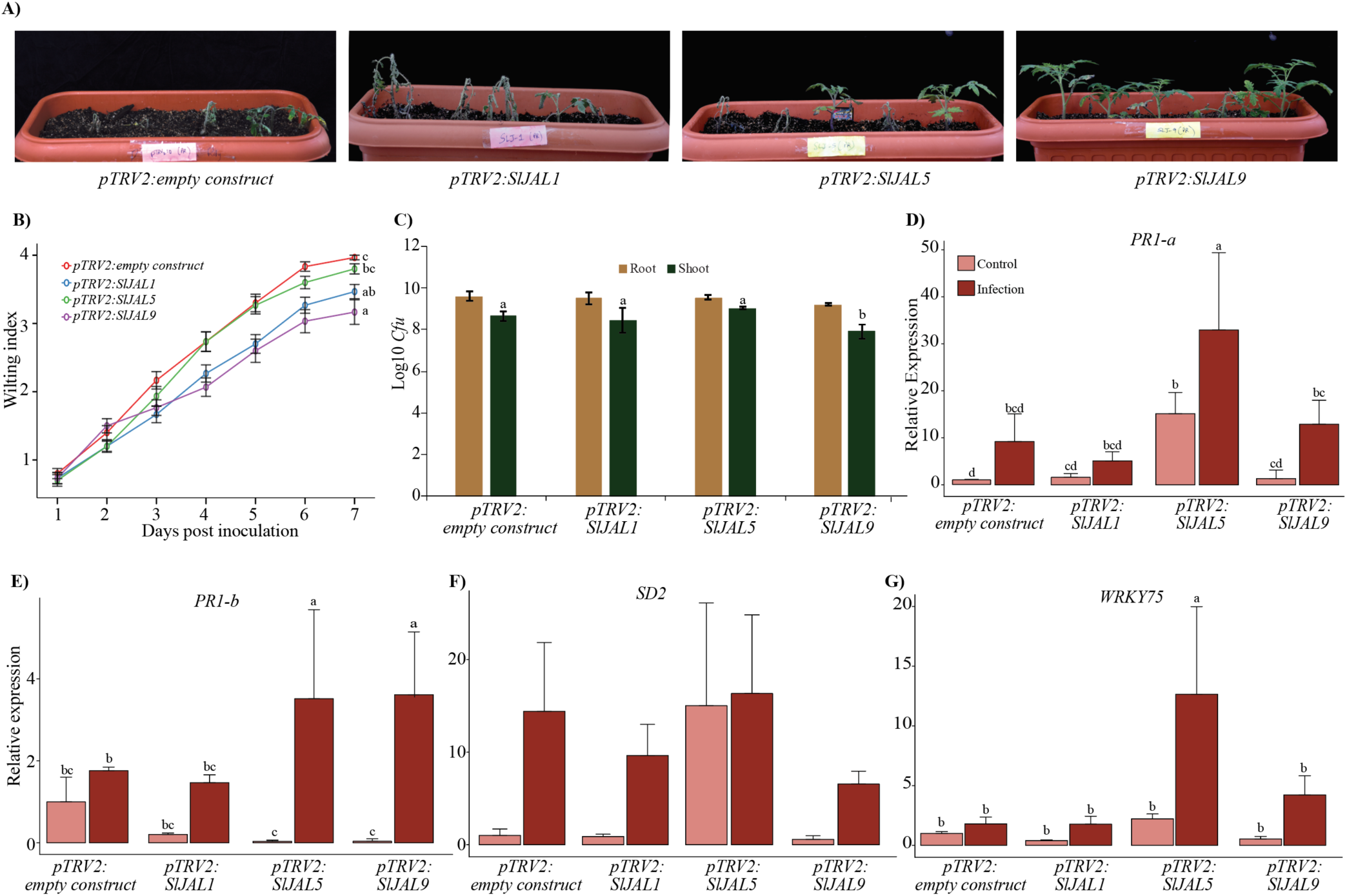
Silencing of *Solanum lycopersicum* JAL orthologs enhances tolerance to *R. pseudosolanacearum*. (A) Representative phenotypes of VIGS lines targeting SlJAL1, SlJAL5, and SlJAL9 following infection with *R. pseudosolanacearum*. (B) Wilting index over time showing delayed disease progression in JAL-silenced plants relative to empty vector controls. Different letters indicate significant differences at 8 dpi. Corresponding statistical details are provided in Supplemental Table S20. (C) Quantification of bacterial colony-forming units in roots and shoots at 7 dpi. (D–G) Relative transcript levels of defense-related genes *PR1A*, *PR1B*, *WRKY75*, and *SD2* in control and JAL-silenced plants. Different letters indicate statistically significant differences (P < 0.05).

## Discussion

### Cytokinin orchestrates root cap immunity at vulnerable entry sites against Ralstonia

*Ralstonia solanacearum* enters primarily through the root system, making the root cap a critical interface between plants and the surrounding soil microbiome. In this region, roots must balance interactions with beneficial microbes while restricting pathogen invasion. The root cap and elongation zone are major sites of auxin–cytokinin interaction and are also highly susceptible to pathogen entry (Chapman and Estelle, 2009; Su et al., 2011). This susceptibility likely reflects their role in accommodating beneficial microbes while maintaining a relatively weaker immune response (Gunawardena and Hawes, 2002; Tsai et al., 2023). As these regions also serve as primary entry points for soil-borne pathogens, localized hormonal regulation is likely to play an important role in controlling microbial access to root tissues. Our study highlights cytokinin signalling as a central regulator of this balance during *Ralstonia* infection. Increased microbial pressure at the root tip was associated with elevated cytokinin signalling, whereas disruption of cytokinin perception strongly compromised immunity, indicating that cytokinin signalling is required for effective defence in vulnerable root tissues (Figure 1). To investigate how cytokinin signalling regulates immune responses in the root cap during pathogen invasion, we focused on the root cap-specific jacalin-associated lectins JAL10 and JAL20. Although plant lectins are widely implicated in stress adaptation and immune regulation, the functions of root cap-associated lectins during root infection remain poorly understood (Esch and Schaffrath, 2017; Witzel et al., 2021). Unlike many previously characterized stress-associated lectins that contain additional signalling or stress-responsive domains, JAL10 and JAL20 belong to the structurally simpler merolectin class (Esch et al., 2022). Their spatially restricted expression in the root cap and rapid repression during *Ralstonia* infection suggest specialized roles in regulating root–microbe interactions (Figure 2).

The spatial repression pattern of JAL10 and JAL20 closely followed cytokinin accumulation in the root tip and was inversely associated with auxin distribution, indicating that local hormone gradients contribute to their regulation. Under physiological conditions, auxin is enriched in the columella root cap, whereas cytokinin predominantly accumulates in the lateral root cap. Upon infection, auxin levels decline while cytokinin levels increase in the root tip (Figure 1), coinciding with repression of JAL10 and JAL20 (Figure 2). This observation is further supported by the positive regulation of JAL expression by auxin (Figure S1), suggesting that infection-induced cytokinin accumulation, together with auxin depletion, contributes to transcriptional repression of these genes.

The presence of ARR-binding motifs within the JAL10 and JAL20 promoter regions, together with ARR1 enrichment near the transcription start site, further supports direct transcriptional repression mediated by cytokinin-responsive type-B transcription factors (Figure 3). Similar transcriptional repression by type-B ARRs has previously been reported for YUCCA genes involved in auxin biosynthesis (Meng et al., 2017), suggesting that cytokinin-mediated transcriptional suppression is not uncommon and represents a broader regulatory mechanism during developmental and stress responses. Importantly, basal cytokinin signalling under physiological conditions does not induce comparable repression of JAL genes, indicating that cytokinin-mediated regulation likely depends on local concentration thresholds and spatial hormone distribution rather than cytokinin presence alone. Together, these findings suggest that cytokinin signalling dynamically remodels root cap transcriptional programs during pathogen attack, enabling rapid transition from a permissive state toward defence activation.

#### Cytokinin-mediated repression of JAL lectins promotes structural and metabolic defence activation

Functional analyses indicate that JAL10 and JAL20 promote pathogen colonization and disease progression. *Loss-of-function jal* mutants delayed *Ralstonia* accumulation, whereas overexpression accelerated infection and disease severity, suggesting that JAL expression contributes to maintaining a permissive root environment under physiological conditions (Figure 4). Their repression during infection may therefore facilitate the transition toward a defence-associated state. This interpretation is consistent with the broader role of the root cap and epidermal cells as a specialized interface that supports microbial colonization while retaining the capacity for rapid immune activation. Since roots continuously interact with diverse beneficial microbes in the rhizosphere, unrestricted activation of defence responses could compromise microbial homeostasis and root function. Dynamic repression of JAL10 and JAL20 during infection may therefore represent a mechanism that allows plants to balance microbial accommodation with inducible defence activation in response to increasing pathogen pressure.

The delayed pathogen progression observed in *jal* mutants suggests that repression of JAL function enhances early defence barriers at the root surface. Given the localization of JAL10 and JAL20 to the endoplasmic reticulum, together with the established roles of ER-associated lectins in protein processing and glycosylation (Lannoo and Van Damme, 2014: Nagashima et al., 2018; Das et al., 2023), JAL proteins may influence the modification or accumulation of proteins required for immune adaptation. This is supported by the pronounced enrichment of proteins associated with cell wall biogenesis and glycan modification observed in *jal* mutants (Figure 5). Notably, the glycoproteome profile of *jal* mutants under basal conditions closely resembled the infection-responsive state of wild-type plants, suggesting that loss of JAL function establishes a preconditioned defence state prior to pathogen challenge. Several enriched proteins, including galactosyltransferases and fucosyltransferases, are associated with cell wall assembly and reinforcement during stress responses (Sarria et al., 2001; Strasser et al., 2007; Zhang et al., 2019) These observations suggest that JAL proteins negatively regulate pathways involved in cell wall remodelling and that their repression during infection enables rapid activation of structural defence programs (Figure 5, Figure S4-S8).

The repression of JAL10 and JAL20 was not restricted to *Ralstonia* infection but was also observed following flg22 treatment and infection with *Fusarium oxysporum* and *Rhizoctonia solani,* two of the most destructive soil-borne fungal pathogens, suggesting that JAL repression forms part of a broader root immune response (Figure S12). Previous studies have shown that pathogen perception activates cytokinin biosynthesis and signalling in roots, including induction of IPT and ARR genes during bacterial and fungal infection (Rich-Griffin et al., 2020; Guo et al., 2021). Together with our findings, this supports a model in which cytokinin-mediated repression of *JAL10 and JAL20* genes represents a general mechanism associated with root immune activation rather than a pathogen-specific response. The enhanced accumulation of cell wall-associated proteins in *jal10 and jal20* mutants is consistent with their delayed pathogen colonization phenotype. Because pathogen invasion is frequently associated with cell wall remodelling and reinforcement, constitutive activation of these pathways likely strengthens the physical barrier against microbial entry.

Interestingly, recent studies have highlighted broader roles for cytokinin signalling in regulating cell wall dynamics during plant development. Cytokinin-mediated signalling has been shown to control secondary cell wall formation by regulating NAC-associated networks through ARR1/10/12-dependent transcription (Didi et al., 2025). Similarly, cytokinin-induced root secondary growth is also initiated by a set of LBD genes in Arabidopsis (Ye et al., 2021). Together with our findings, these studies suggest that cytokinin functions as a broader regulator of cell wall plasticity, dynamically coordinating wall-associated programs in both developmental and immune contexts. While cytokinin suppresses premature secondary wall formation during development, our results suggest that pathogen-induced cytokinin signalling promotes defence-associated cell wall remodelling by repressing JAL lectins, highlighting the context-dependent regulation of cell wall dynamics during stress adaptation.

In addition to changes in glycoprotein accumulation, *jal* mutants displayed enhanced phenylpropanoid-associated metabolism. The phenylpropanoid pathway generates a diverse range of defence-associated metabolites and also contributes to lignin biosynthesis, an important component of cell wall reinforcement (Figure 6). Increased abundance of UGT-associated proteins, together with elevated levels of phenylpropanoid metabolites, suggests coordinated activation of both structural and metabolic defence pathways in *jal* mutants. Lignification is closely associated with reactive oxygen species production and strengthening of the cell wall during immune responses (König, S., et al, 2014; Pascual, et al, 2016). Consistent with this, enhanced expression of *RBOHD* in *jal10 and jal20* mutants likely leads to elevated levels of ROS and thereby enhanced activation of lignin-associated defence processes. The simultaneous enrichment of cell wall-associated proteins, phenylpropanoid metabolites, and ROS-related responses suggests that *loss-of-function* of *jal10 and jal20* promotes a constitutively primed defence state that strengthens both physical and biochemical barriers against pathogen invasion.

#### Tomato JAL lectins contribute to conserved regulation of root susceptibility *to Ralstonia*

The identification of structurally related JAL proteins in tomato and the reduced susceptibility of *SlJAL*-silenced plants further indicate that JAL proteins’ immune regulatory role is conserved across species. Although tomato contains fewer JAL homologs than Arabidopsis, silencing of specific *SlJAL* genes was sufficient to reduce disease susceptibility, suggesting a conserved contribution to pathogen colonization and disease progression in roots (Figure 7, Figure S10).

Collectively, our findings support a model in which pathogen invasion at the root tip triggers activation of cytokinin signalling, leading to transcriptional repression of the root cap-specific lectins JAL10 and JAL20 through ARR-mediated regulation. Under physiological conditions, JAL proteins contribute to maintaining a permissive root environment that supports microbial interactions. Upon infection, however, cytokinin-dependent repression of JAL expression promotes activation of cell wall remodelling, glycan modification, phenylpropanoid metabolism, and ROS-associated defence pathways (Figure 8). These coordinated structural and metabolic responses reinforce the root surface barrier, restrict pathogen colonization, and enhance root immunity. The conservation of JAL-mediated susceptibility in tomato further suggests that this cytokinin–JAL regulatory module represents a broader mechanism by which plants dynamically regulate microbial interactions at the root–soil interface and adapt to soil-borne pathogen attack.

**Figure 8.**
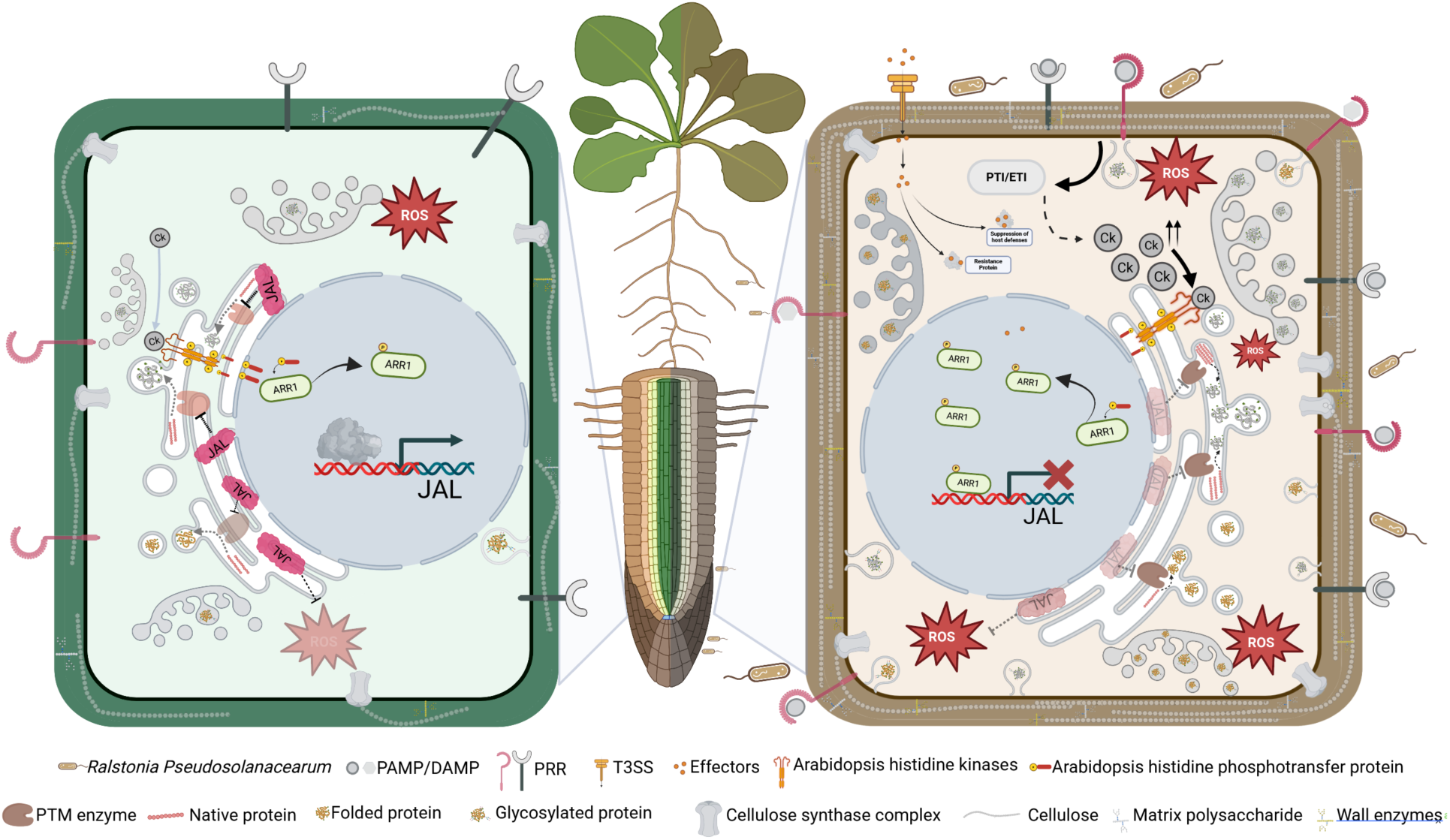
Proposed model illustrating cytokinin-mediated repression of JAL lectins enables cell wall reinforcement and restricts *Ralstonia* invasion at the root tip. Under basal conditions, JAL10 and JAL20 are expressed in the root cap and may contribute to maintaining a permissive environment for root–microbe interactions. Upon pathogen invasion and increasing microbial load, cytokinin biosynthesis and signalling are enhanced in the root tip. Activated type-B ARR transcription factors directly repress JAL10 and JAL20 expression, leading to activation of cell wall biogenesis and phenylpropanoid-associated defense pathways. This results in increased accumulation of cell wall-associated glycoproteins, lignin precursors, flavonoids, and other defense-related metabolites, ultimately reinforcing structural and chemical barriers that restrict Ralstonia invasion and systemic colonization.

## Acknowledgments

We acknowledge Prof. Suvendra Kumar Ray (Tezpur University) for providing the *Ralstonia pseudosolanacearum F1C1* strain. We thank Dr. Gopaljee Jha (NIPGR) for assistance with VIGS experiments. We also acknowledge Dr. Satish Mutyam for support with metabolomics. Technical assistance from Rama Sai Venkata Marthi (Project Associate) and Anurag Kumar (BSMS, IISER Mohali) is sincerely appreciated.

## Author contributions

E.R. conceived the project, secured funding, and supervised the study. A.P.G. performed the experiments and conducted data analysis. V.K. and S.B.S carried out the ChIP–qPCR experiments. A.D. and K.A. performed LC-MS and processed the glycoproteomics data. K.K.D, A.P.G, and S.H generated the *JAL10* and *JAL20* functional mutants. A.P.G and I.T.K generated the VIGS constructs. A.P.G. and E.R. wrote the original draft, and all authors contributed to reviewing and editing and approved the final version.

## Data availability

The mass spectrometry proteomics data have been deposited to the ProteomeXchange Consortium via the PRIDE partner repository with the dataset identifier PXD073654.

## Supporting Information (SI)

Supplemental Methods

Supplemental Figures S1-S12

Supplemental Tables S1-S21

**Figure S1.).**
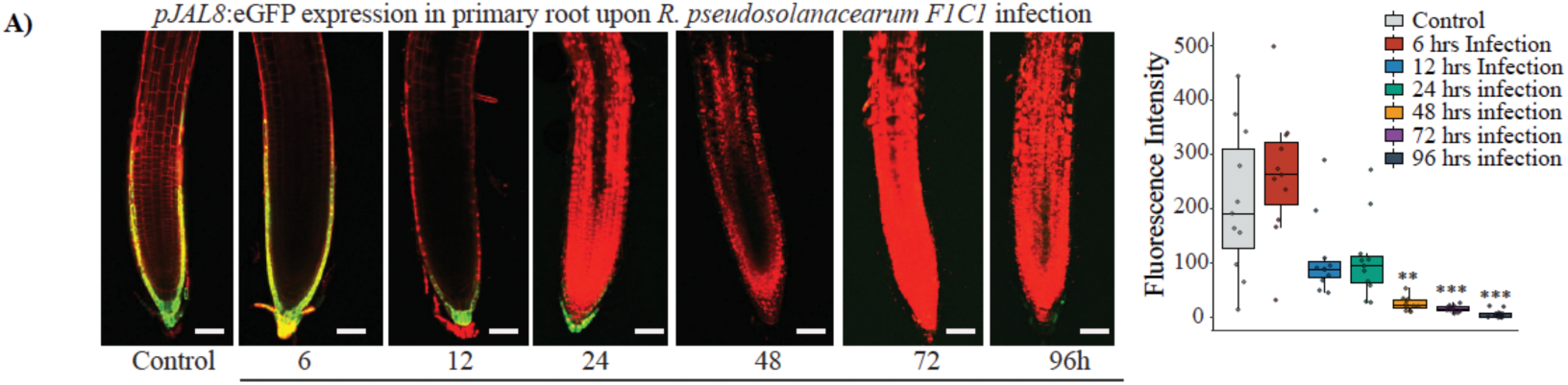
JAL family members are repressed by pathogen-associated signals. (A) Confocal images showing *pJAL8* reporter activity during infection with *R. pseudosolanacearum* over a 96-hour time course. GFP fluorescence is shown in green. Scale bar, 100 μm.

**Figure S2.).**
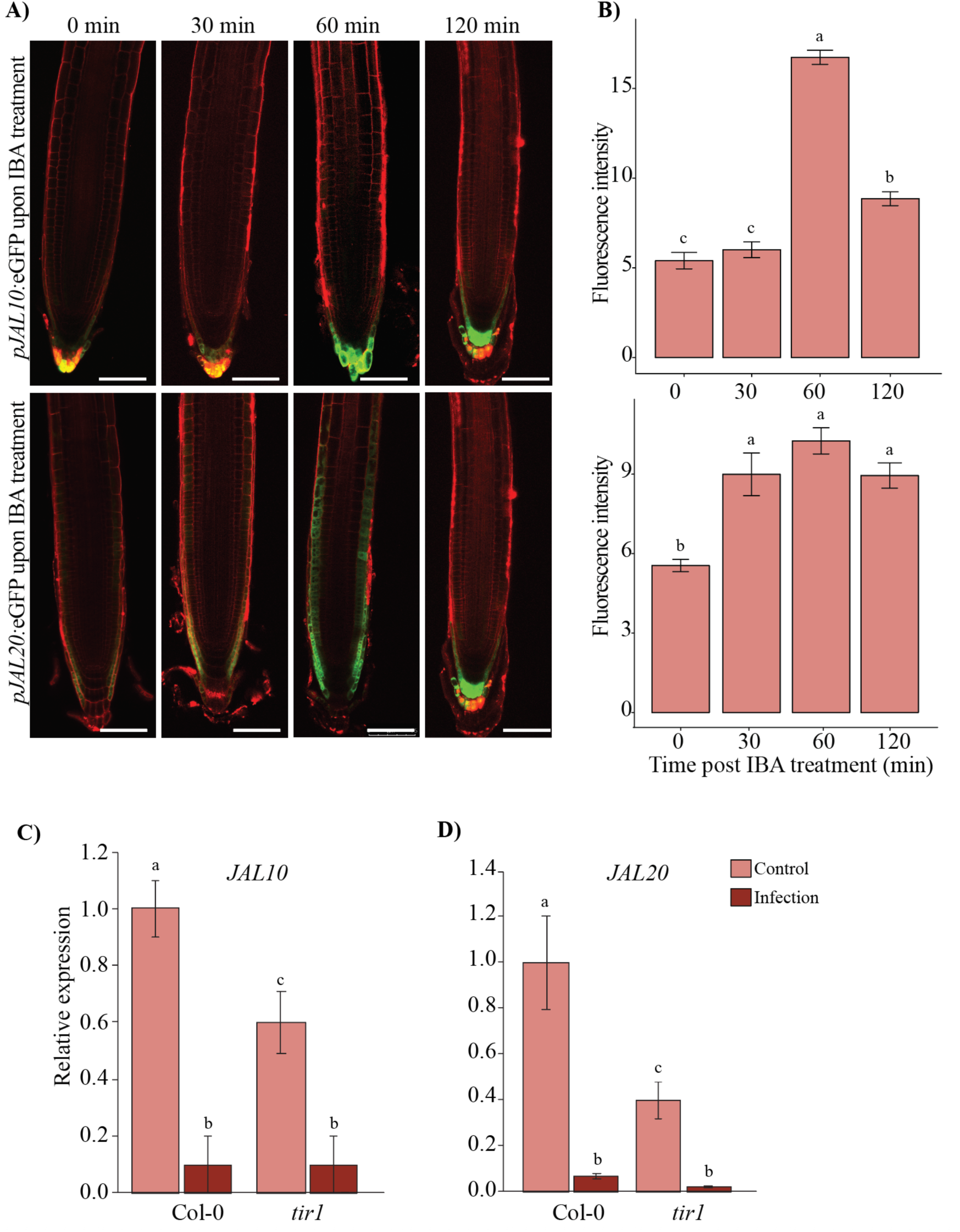
Auxin positively regulates JAL10 and JAL20 expression. (A) Confocal images showing pJAL10 and pJAL20 reporter activity following treatment with 1 μM IBA. GFP fluorescence is shown in green and cell walls stained with propidium iodide in red. (B) Quantification of fluorescence intensity following IBA treatment. Data represent three independent biological replicates (n = 15 per experiment). (C, D) Relative transcript levels of *JAL10* and *JAL20* in the auxin receptor mutant *tir1* under control and infection conditions. Statistical significance was determined using two-way ANOVA followed by Fisher’s LSD test.

**Figure S3.).**
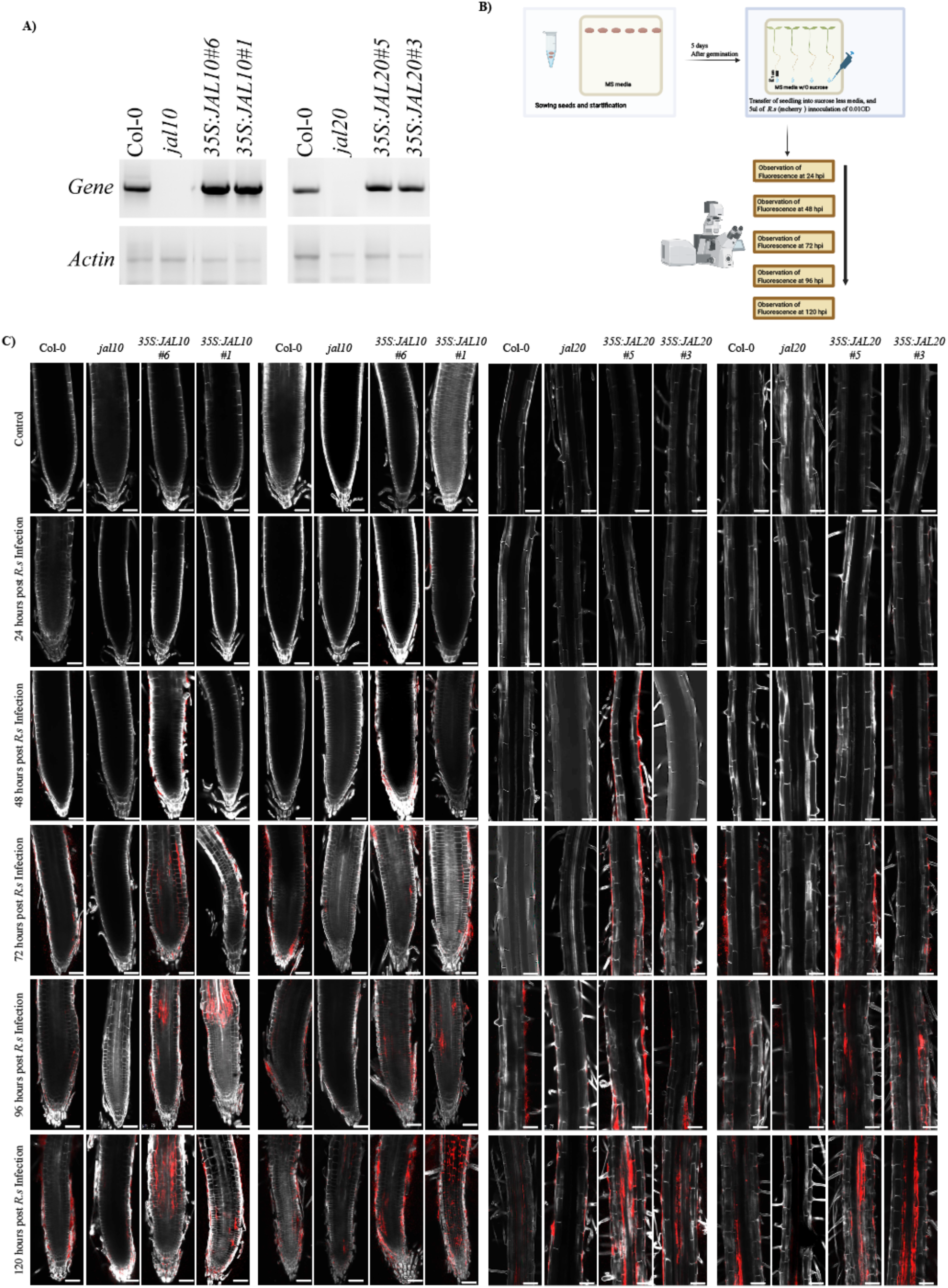
Validation of JAL functional mutants and invasion assay experimental setup. (A) Semi-quantitative RT–PCR analysis of *JAL10* and *JAL20* expression in Col-0, knockout mutants, and overexpression lines. *ACTIN* was used as an internal control. (B) Schematic representation of the *in vitro* invasion assay. (C) Representative confocal images showing progression of *R. pseudosolanacearum-mCherry* during invasion assays from 24 to 120 hpi in JAL10 and JAL20 functional mutants. Bacterial fluorescence is shown in red and cell walls stained with calcofluor white. Scale bar, 50 μm.

**Figure S4.).**
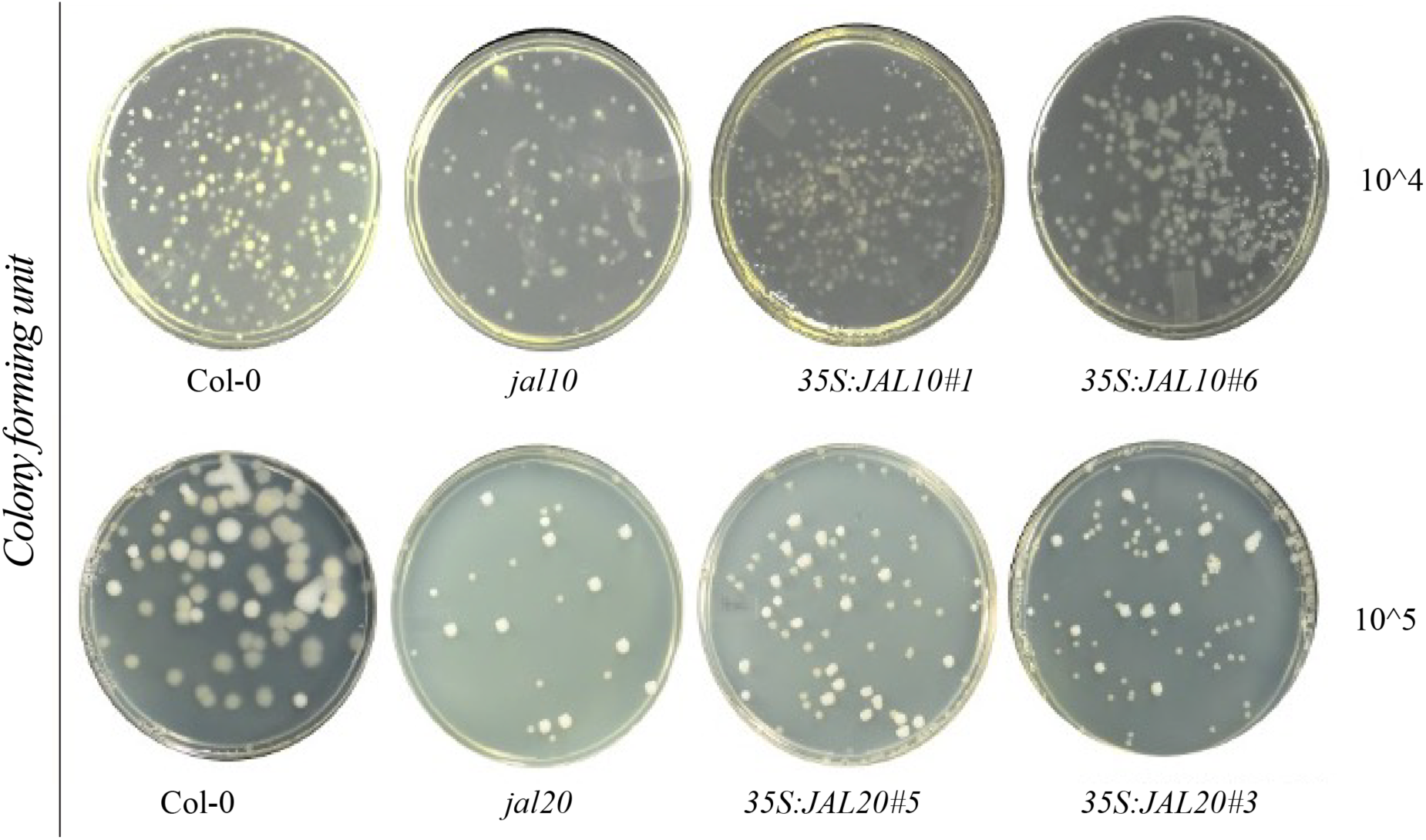
Representative images of bacterial colonies formed on BG agar plates following CFU assays in JAL10 and JAL20 functional mutants.

**Figure S5.).**
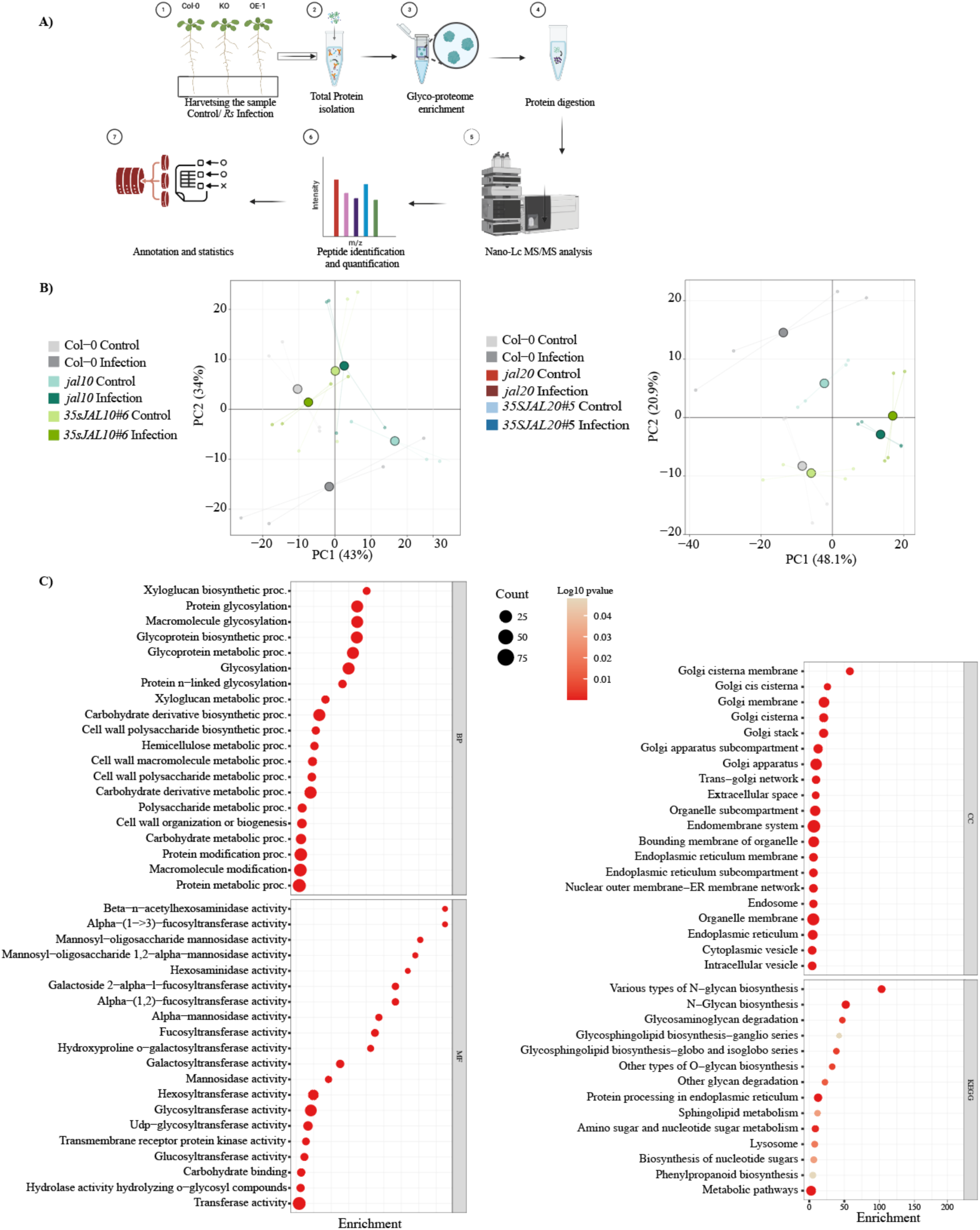
Overview of glycoproteomic analysis. (A) Schematic workflow of the glycoproteomic analysis. (B) Principal component analysis of glycoproteomic profiles from JAL10 and JAL20 functional mutants. (C) Gene Ontology enrichment analysis of the total glycoproteome, including biological process (BP), molecular function (MF), cellular component (CC), and KEGG pathway enrichment.

**Figure S6.).**
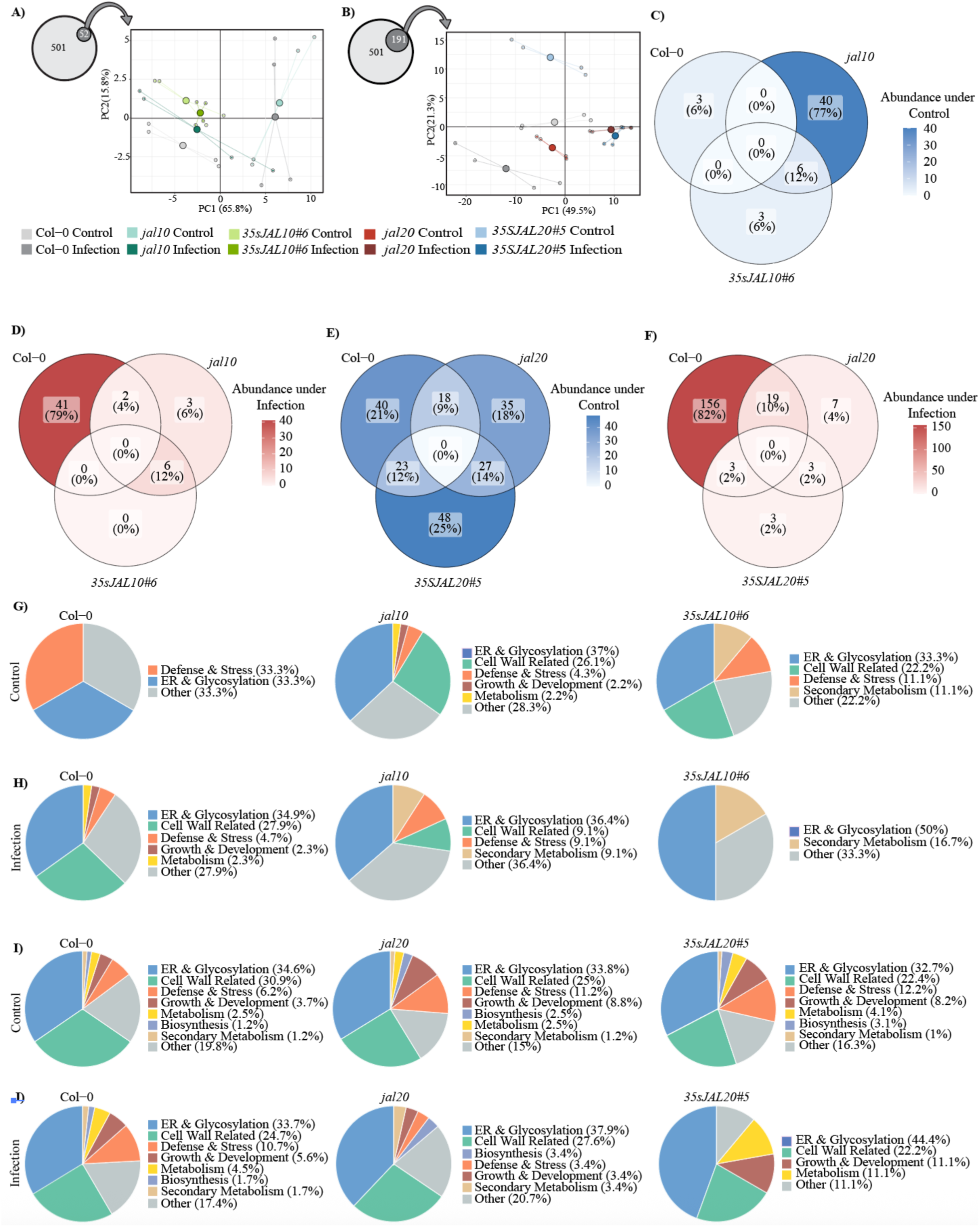
Distribution of significantly altered glycoproteins in JAL mutants. (A, B) Principal component analysis of significantly altered proteins in JAL10 and JAL20 functional mutants.(C-F) Relative abundance of significantly altered proteins under control and infection conditions. (G-J) Distribution of enriched biological processes in JAL10 and JAL20 functional mutants and Col-0 under control and infection conditions.

**Figure S7.).**
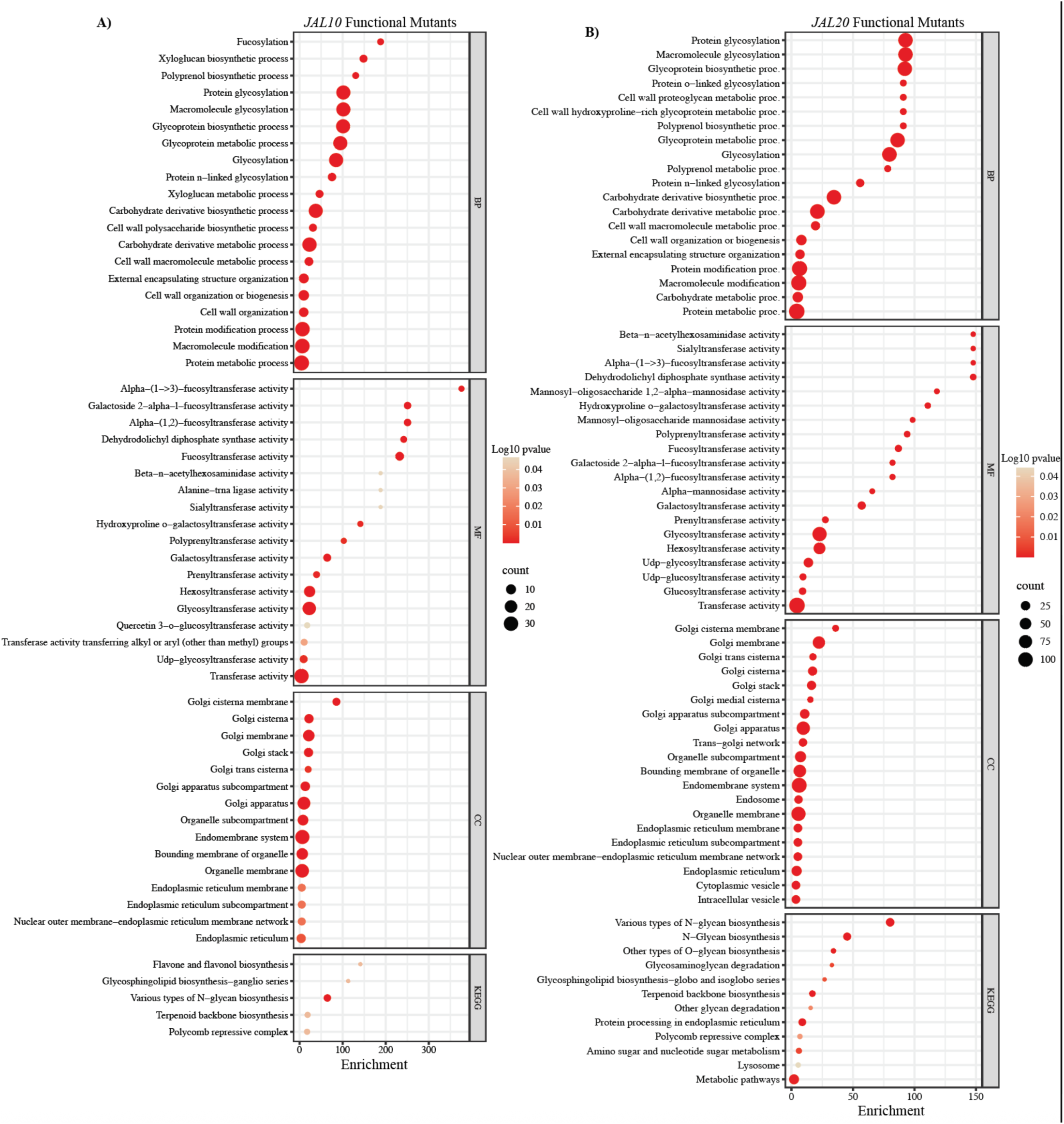
Functional enrichment analysis of significantly altered proteins. Gene Ontology enrichment analysis of significantly altered proteins in JAL10 and JAL20 functional mutants showing enriched biological process (BP), molecular function (MF), cellular component (CC), and KEGG pathway categories.

**Figure S8.).**
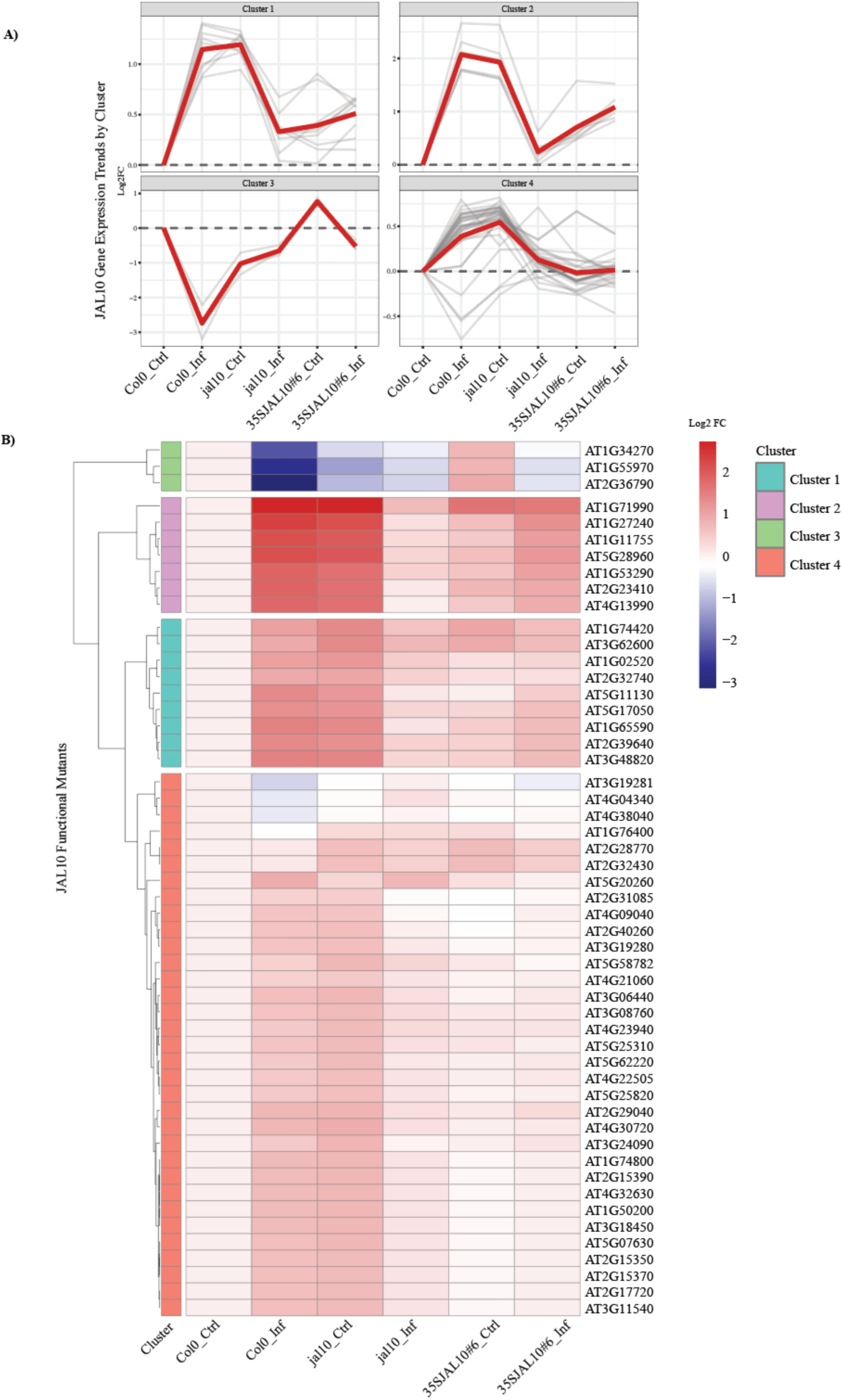
Clustering analysis of glycoproteins in JAL10 functional mutants. (A) Expression patterns of significantly altered proteins in JAL10 functional mutants under control and *R. pseudosolanacearum* infection conditions. (B) Heatmap showing K-means clustering of significantly altered proteins relative to Col-0 control conditions.

**Figure S9.).**
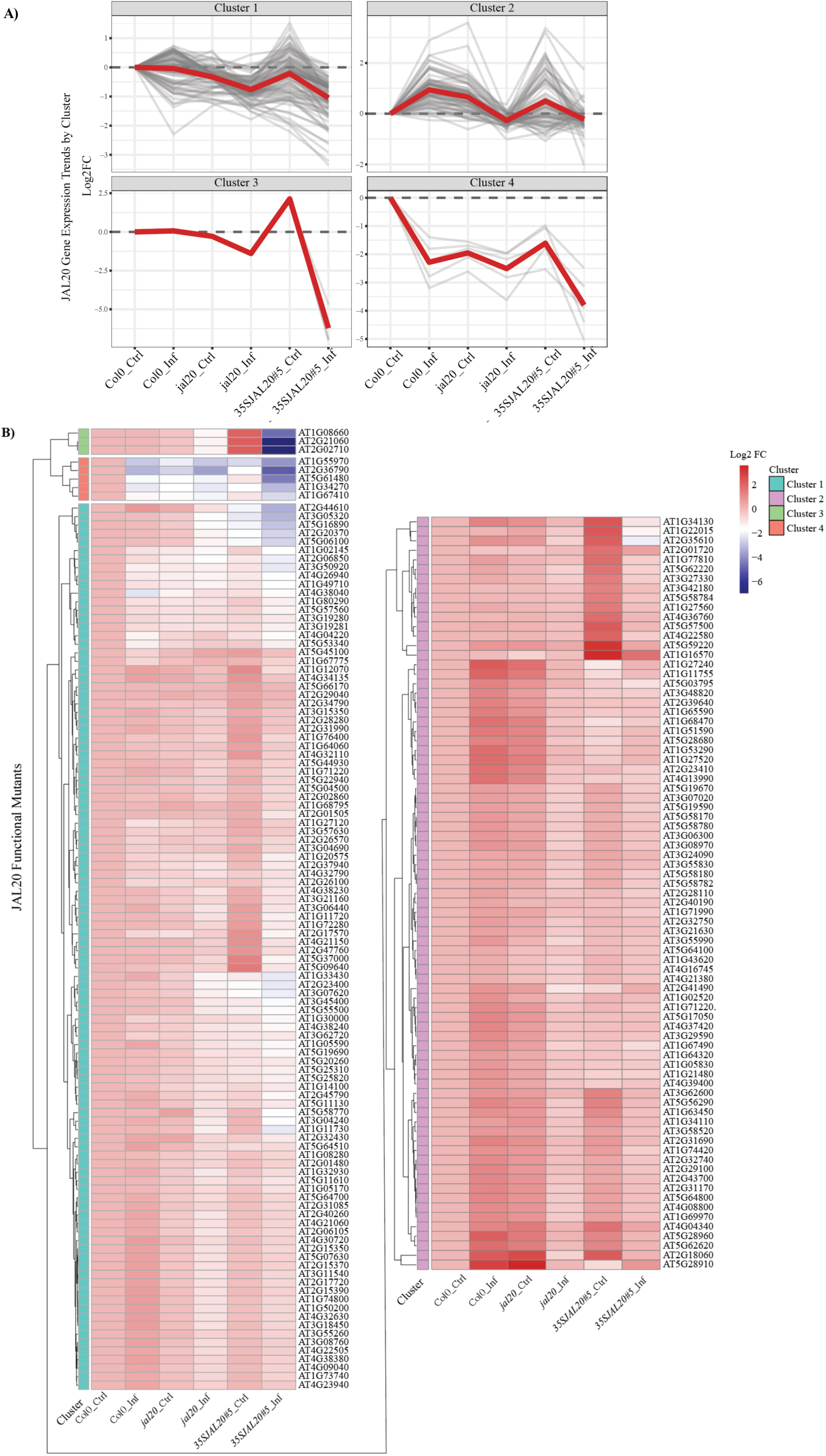
Clustering analysis of glycoproteins in JAL20 functional mutants. (A) Expression trends of significantly altered proteins in JAL20 functional mutants under control and R. pseudosolanacearum infection conditions. (B) Heatmap showing K-means clustering of significantly altered proteins relative to Col-0 control conditions.

**Figure S10.).**
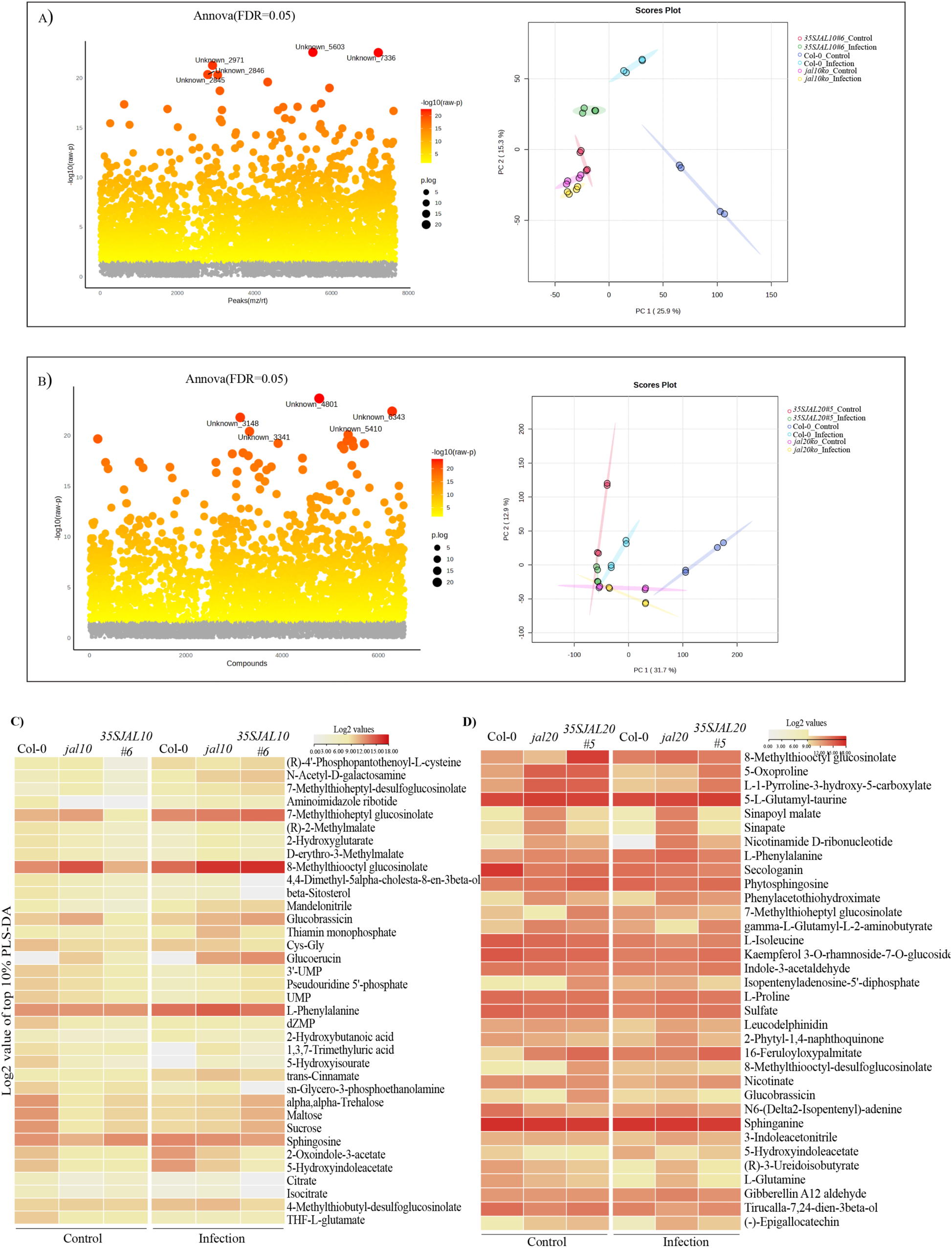
Metabolomic profiling of JAL functional mutants. (A, B) Principal component analysis and distribution of significantly altered metabolites identified using one-way ANOVA with FDR correction (FDR ≤ 0.05). Analyses were performed using MetaboAnalyst. (C, D) Heatmaps showing the top 10% of metabolites identified by PLS-DA analysis in JAL10 and JAL20 functional mutants.

**Figure S11.).**
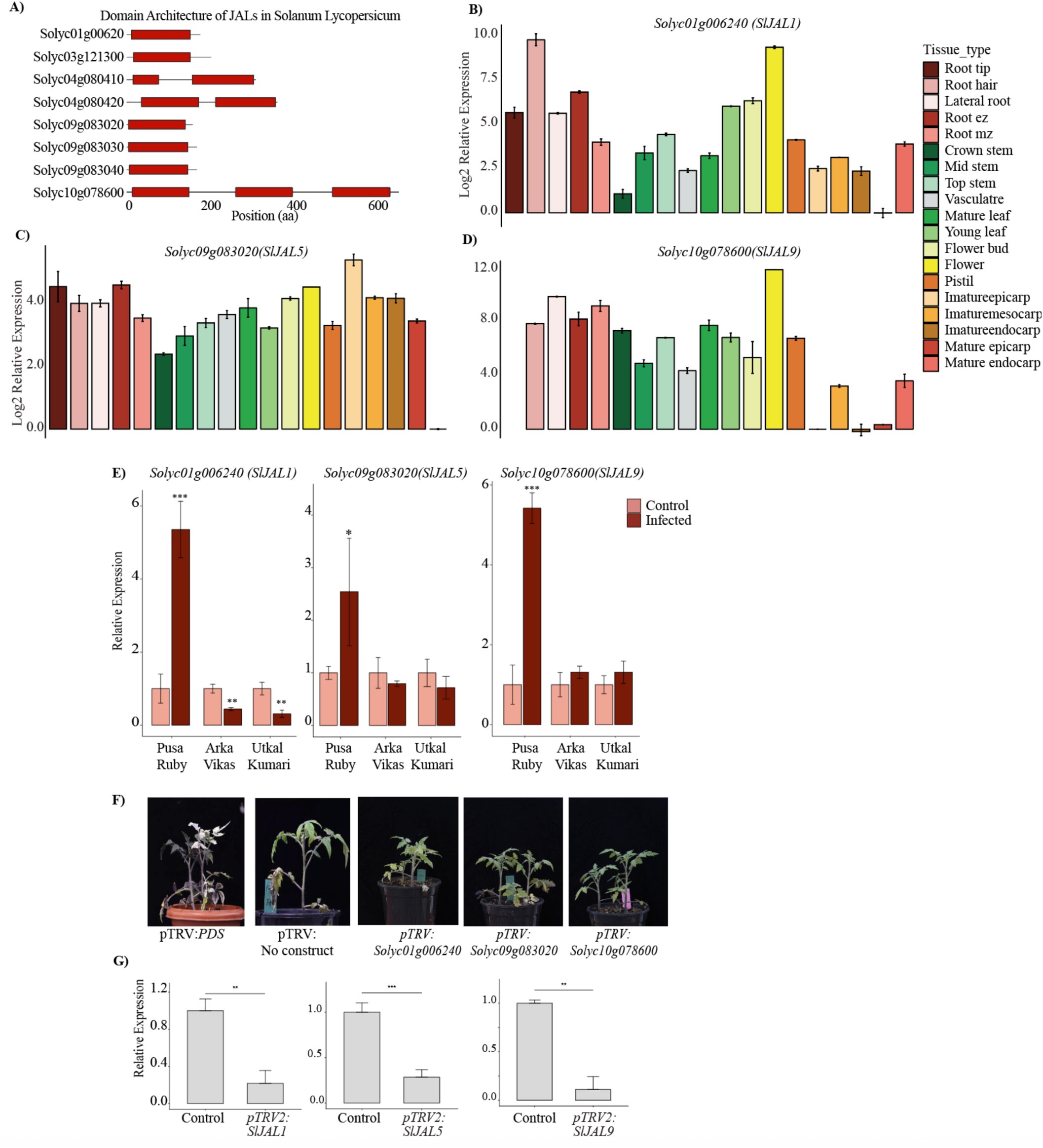
Identification and functional analysis of tomato JAL orthologs. (A) Domain architecture of tomato jacalin-associated lectins identified in *Solanum lycopersicum*. (B-D) Tissue-specific transcript abundance of SlJAL1, SlJAL5, and SlJAL9. (E) Relative transcript abundance of *SlJAL1*, *SlJAL5*, and *SlJAL9* following *R. pseudosolanacearum* infection in tomato varieties with differing susceptibility levels. in the Pusa Ruby- highly susceptible variety, Arka Vikas- moderately susceptible variety, and Utkal Kumari- tolerant variety (F) Phenotype of pTRV plants used as a positive control for VIGS efficiency. (G) Validation of transcript silencing in SlJAL1-, SlJAL5-, and SlJAL9-silenced plants. Statistical significance was determined using Student’s *t*-test (*P ≤ 0.05, **P ≤ 0.01, ***P ≤ 0.001).

**Figure S12.).**
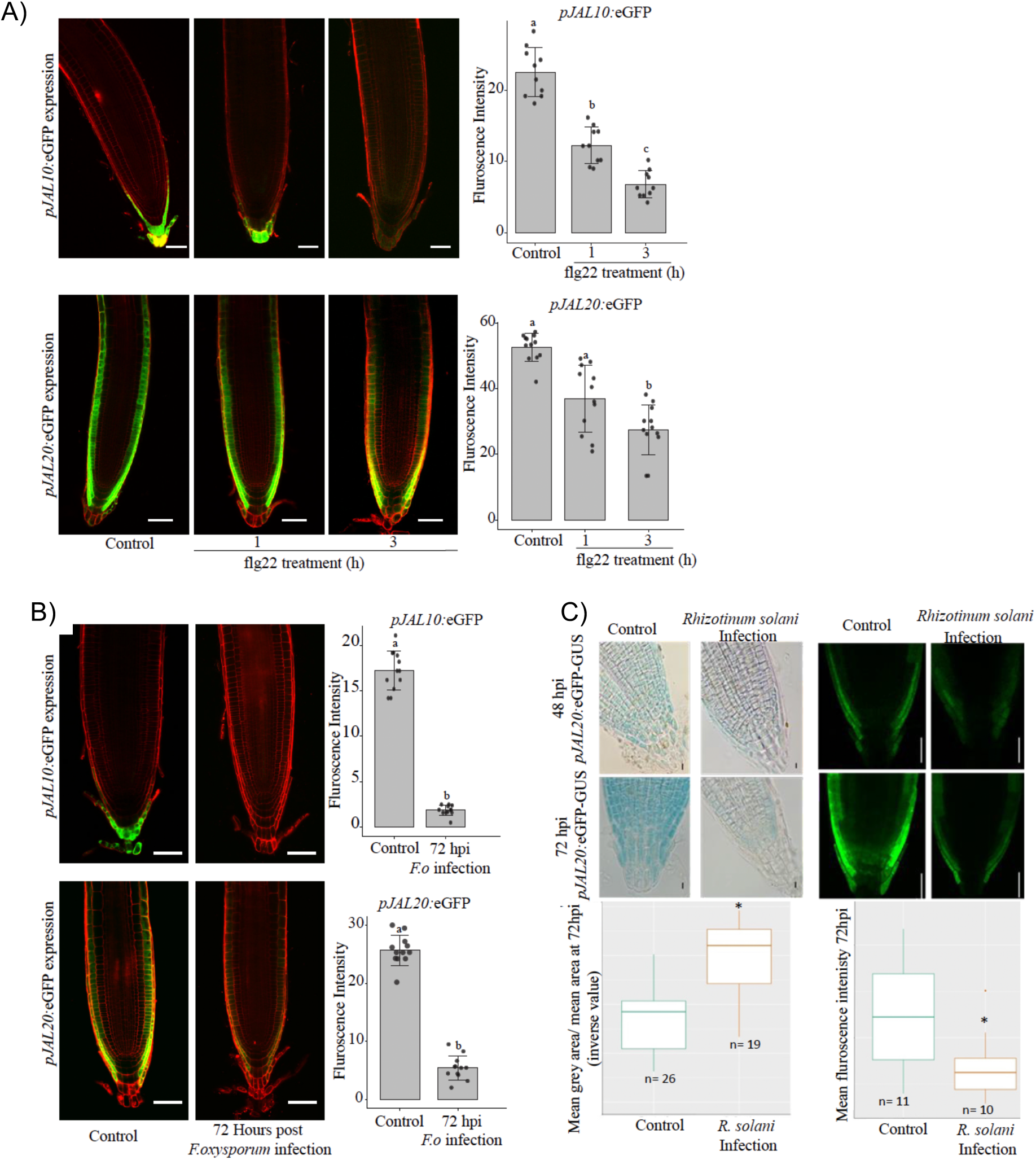
JAL family members are repressed by pathogen-associated signals. (A) Reporter activity of *pJAL10* and *pJAL20* following flg22 treatment for 3 h. Scale bar, 75 μm. (B) Reporter activity of *pJAL10* and *pJAL20* following *Fusarium oxysporum* infection at 72 hpi. Scale bar, 75 μm. (C) Reporter activity of pJAL20 following *Rhizoctonia solani* infection. Bright-field scale bar, 10 μm; confocal scale bar, 90 μm. Statistical significance was determined using the Kruskal–Wallis test followed by Dunn’s multiple comparison test (*P ≤ 0.05, **P ≤ 0.01, ***P ≤ 0.001). Three independent biological replicates were performed (n = 15 per experiment).

