## Supplemental Information for "Cytokinin-mediated repression of jacalin lectins reinforces root immunity"

**Supplemental Methods**

**Infection assays**

For infection assays, *Ralstonia pseudosolanacearum F1C1*, obtained from S.K. Kumar Ray and cultured in BG broth (10 g peptone, 1 g yeast extract, 1 g casamino acid), was used as previously described by Kumar et al. (2017). The bacterial culture was revived from glycerol stocks and grown in BG broth. For infection experiments, a secondary culture was prepared and harvested by centrifugation at 5,000 × g for 10 min. The resulting bacterial pellet was washed twice with autoclaved distilled water to remove residual media components. Subsequently, the cells were resuspended in sterile distilled water, and the bacterial suspension was adjusted to the desired optical density at 600 nm (OD₆₀₀) to ensure consistent inoculum concentration across experiments.

For analysis of promoter activity in response to *R. pseudosolanacearum*, 5-day-old seedlings were transferred to half-strength MS plates lacking sucrose and MES. A secondary culture of *R. pseudosolanacearum*-mCherry was prepared, and the bacterial suspension was adjusted to an OD₆₀₀ of 0.01. The inoculum was then applied directly to the root tip of 5ul. Following inoculation, plates were incubated at 28 °C to allow infection progression. Promoter activity was monitored at the indicated time points using confocal laser scanning microscopy, employing Leica SP8 and Olympus FV3000 systems.

**In vitro invasion assay**

To determine pathogen invasion in roots, Arabidopsis thaliana seedlings Col-0, *jal10, jal20, 35S:JAL10#6, 35S:JAL10#1, 35S:JAL20#5,* and *35S:JAL20#6* were used. Five-day-old seedlings were transferred to half-strength MS plates lacking sucrose and MES. A secondary culture of *R. pseudosolanacearum*-mCherry was prepared, and the bacterial suspension was adjusted to an OD₆₀₀ of 0.01. The inoculum was applied approximately 1 cm away from the root tip to monitor directional invasion toward the root (Supplemental file S4B).

Following inoculation, plates were incubated at 28 °C to allow infection progression. Imaging was performed using a Leica SP8 confocal laser scanning microscope at 24-hour intervals, up to 120 hours post-inoculation, to track the dynamics of pathogen invasion.

***CFU assay***

To determine bacterial colonization, *Arabidopsis thaliana* lines *Col-0, jal10, jal20, 35S:JAL10#6, 35S:JAL10#1, 35S:JAL20#5, and 35S:JAL20#6, ahk2/ahk4, pPYK10:CKX3* were subjected to colony-forming unit (CFU) assays following the method described by Yu, Gang, et al. (2023). In brief, 5-day-old seedlings were transferred to half-strength MS plates lacking sucrose and MES. A secondary culture of *R. pseudosolanacearum*-mCherry was adjusted to an OD₆₀₀ of 0.01 and inoculated directly onto the root tip. Plates were then incubated at 28 °C for 3 days. Following infection, roots and shoots were carefully harvested, washed to remove surface-associated bacteria, and dried prior to weighing separately. Each sample was homogenized in 200 µL of sterile water using a Retsch mill at 30 cycles for 2 minutes, and the volume was subsequently adjusted to 1,000 µL. An aliquot of 20 µL of the homogenate was then diluted in 180 µL of sterile water, followed by serial dilutions up to 10⁻⁵. A 100 µL aliquot from the 10⁻⁵ dilution was spread onto BG agar plates supplemented with 30 µg/mL gentamycin and incubated at 28 °C. Colonies of *R. pseudosolanacearum* were visible after 36 hours, counted, and quantified as CFU per milliliter. The experiment was conducted with six biological replicates.

***Soil-Drench assay***

Five-day-old germinated seedlings *Col-0, jal10, jal20, 35S:JAL10#6, 35S:JAL10#1, 35S:JAL20#5, and 35S:JAL20#6, ahk2/ahk4, pPYK10:CKX3* were transferred to soil and grown under short-day conditions for up to 3 weeks. For infection, a drench assay was performed using a secondary culture of *R. pseudosolanacearum*, adjusted to an OD₆₀₀ of 0.1. The bacterial suspension was applied by soil drenching, and plants were subsequently maintained in a growth chamber at 28 °C. Disease progression was monitored by assessing wilting symptoms post-inoculation, and representative images were captured using a DSLR camera. Each experiment included at least 15 infected plants and was independently replicated. The wilting index was determined following the method described by Hiles et al. (2024). Briefly, disease severity was scored on a scale from 0 to 4: 0 = no visible symptoms; 1 = ≤25% wilting; 2 = ≤50% wilting; 3 = ≤75% wilting; and 4 = ≤100% wilting of leaves. Based on these scores, the disease index (DI) was calculated using the formula: DI = Nw/N, where Nw represents the number of wilted leaves and N represents the total number of leaves.

**Glycoprotemics**

***Protein Extraction, Processing and Peptide preparation***

Arabidopsis seedlings Col-0, *jal10, jal20, 35S:JAL10#6, 35S:JAL20#5*, were grown in Murashige and Skoog medium (MS) media for a duration of five days. Subsequently, they were transferred to a sucrose less, reduced media and inoculated with *R. pseudosolanacearum* along mock treatment as control. After a period of 48 hours of infection, the root tips of the seedlings, measuring up to 0.5 cm in length, were excised and subsequently collected using snap freezing. Protein samples were prepared according to the protocol mentioned in Pierce Plant Total Protein Extraction Kit as mentioned in the kit (Thermo Fisher Scientific, Rockford, Illinois, USA). The concentration of the enriched glycoprotein was analysed using the BCA kit (Thermo Fisher Scientific, Rockford, Illinois, USA). The enriched glycoproteins were redissolved in 6 M urea, 50 mM ammonium biocarbonate buffer. The protein samples were treated with DTT for 1 hour at 37°C for 1 hour followed by IAA addition for 30 minutes in dark at room temperature. Ammonium biocarbonate buffer was further added to reduce the urea concentration to 0.6 M before digestion. For digestion, trypsin (Promega, Madison, WI, USA) was added at 1:25 ratio for 16 hours at 37°C. Formic acid (1%) was used to stop the reaction and the peptides were desalted using Pierce™ C18 Spin Columns (Thermo Fisher Scientific, Rockford, Illinois, USA). The peptides were dried using speedy-Vac and reconstituted in 3% ACN in 0.1% formic acid.

***Protein Identification by Mass Spectrometry***

Peptide samples were injected onto nanoEase M/Z Symmetry C18 trap Column (100 Å, 5 μm, 180 μm × 20 mm) (Waters, Milford, USA), followed by a reversed-phase nanoEase M/Z HSS T3 Column (100 Å, 1.8 μm, 75 μm × 100 mm) (Waters, Milford, USA). MS solvents e.g., 0.1% formic acid (A) and acetonitrile with 0.1% formic acid (B) were used for chromatographic separation of glycopeptides. Peptide mixtures were run in Waters LC/MS G2-XS Q-TOF (Milford, Massachusetts, USA) at a flow rate of 300 nL/min. A gradient of 120 min was performed for the sample analysis, as described (time in minutes, %B): 5 min, 5%; 95 min, 40%; 100 min, 85%; 120 min, 5%. Sample flow rate was 300 nL/min with MS operating parameters as follows: positive polarity, sensitivity mode, scan time 0.5 seconds, continuum data format, and collision energy ramp from 15 to 40. While the sampling cone and capillary voltage were set at 40 V and 3 kV respectively to ensure stable spray formation. Leucine enkephalin (m/z = 556.2771), the lock mass calibrant, was infused at a flow rate of 300 nL/min, with acquisition at 30-sec intervals to ensure the precision of the mass measurement.

***Database search and peptide identification***

Mass spectrometry (MS) data were acquired using Masslynx v4.2 (Waters, Milford, USA) to generate raw files. MS raw files were processed using the ion accounting algorithm within the Progenesis QIP (Nonlinear Dynamics, UK) search engine against the UniProt Arabidopsis thaliana (Mouse-ear cress) to obtain peptide and protein identifications. The threshold for both protein and peptide identification was 0.01 false discovery rate (FDR), a maximum of 1 missed cleavage with trypsin as the enzyme, a precursor mass tolerance of 20 ppm, and a fragment mass tolerance of 40 ppm. Carbamidomethyl (C) was selected as fixed modification while C-Mannosyl (W), Glycation (N terminal), O-GlcNac (ST), oxidation (M) options were selected as variable modifications. Utilizing the Progenesis QIP quantitation approach, which aligns detectable characteristics of all runs for relative quantitative analysis, the identified proteins were quantified. When at least two different peptides were detected in MS/MS, protein identifications were considered as confident.

***Glycoprotemics analysis***

Gene Ontology (GO) enrichment analysis was performed using ShinyGO 0.85 (Ge et al., 2020), and the top enriched categories were visualized using SR plots (Tang et al., 2023). Pairwise comparisons among knockout, overexpression, and Col-0 lines were conducted using one-to-one t-tests for each condition. Statistical significance was determined using a Benjamini–Hochberg false discovery rate (FDR) threshold of 0.05. Proteins with a fold change ≥1.5 were classified as upregulated, whereas those with a fold change ≤0.6 were considered downregulated. Proteins meeting these criteria in any comparison were retained for downstream analyses to capture condition-dependent changes in the glycoproteome. Application of these criteria resulted in the identification of 52 and 191 proteins in *JAL10* and *JAL20* functional mutants, respectively, compared with Col-0. Protein annotation was performed by mapping UniProt IDs obtained from the analysis to Arabidopsis thaliana proteins using UniProt BLAST, selecting the closest homologs (January 2026)

For quantitative analysis, raw protein abundance values from biological replicates were averaged for each genotype (Col-0, knockout, overexpression) under control and infection conditions, separately for *JAL10* and *JAL20*. To identify relatively abundant proteins within each treatment group, row-wise Z-scores were calculated independently for control and infection datasets across the three genotypes. The Z-score is a standardized measure that indicates how far a given value deviates from the mean, calculated as (X − μ) / σ, where X represents the raw value, μ is the mean, and σ is the standard deviation. A Z-score > 0 indicates that the value is above the mean, a Z-score = 0 indicates that it is equal to the mean, and a Z-score < 0 indicates that it is below the mean. Proteins with positive Z-scores (>0) were considered to exhibit above-average abundance and were classified as “highly abundant” within that specific genotype and condition. These assignments were further used for Venn diagram representation, where proteins were allocated to a genotype if their Z-score exceeded >0 (Kryuchkova-Mostacci, N., & Robinson-Rechavi, M. ,2017, Eisen, M. B., et al, 1998).

Functional categorization of proteins was initially for selected proteins were obtained using ShinyGO 0.85 and subsequently manually curated into broader functional groups for pie chart, as detailed in Supplemental Table 11 & 13. Log₂ fold changes were calculated relative to the Col-0 control condition.

**Metabolomics**

Arabidopsis seedlings Col-0, *jal10, jal20, 35S:JAL10#6, 35S:JAL20#5* were grown in Murashige and Skoog medium (MS) media for a duration of five days. Subsequently, they were transferred to a sucrose less, reduced media and inoculated with Ralstonia along mock treatment as control. After a period of 48 hours of infection, the root tips of the seedlings, measuring up to 0.5 cm in length, were excised and subsequently collected using snap freezing. Further the root sample were freeze dried, and then dissolved in roots were homogenized in molecular grade methanol containing 0.1 formic acid, and 0.5 mM lidocaine (internal control) and subjected to LCMS.

The LC–MS system consisted of an Agilent 1290 Infinity II liquid chromatography system (Agilent Technologies, Santa Clara, CA, USA) equipped with a quaternary pump (G7104A), a multisampler (G7129B), and a multicolumn thermostat (G7116B), hyphenated to an Agilent 6545XT Q-TOF mass spectrometer. Chromatographic separation was carried out on a C18 column at 40.0°C with a mobile phase flow rate of 0.400 ml min⁻¹. The elution was conducted using Water (Solvent A) and Acetonitrile (Solvent C) in the following gradient: 0–0.5 min maintained at 2% C; 0.5–5.0 min linear ramp from 2% to 98% C; 5.0–6.0 min maintained at 98% C; 6.0–6.1 min return to 2% C; and 6.1–8.0 min re-equilibration at 2% C. The samples (5.00 µl) were injected into the inlet port with the sampler thermostat maintained at 4°C. The mass spectrometer was operated in positive and negative ionization modes using a Dual Agilent Jet Stream ESI source with the following settings: capillary voltage of 3.5 kV (3500 V), nozzle voltage of 1.0 kV (1000 V), drying gas temperature of 300°C at a flow rate of 8 l min⁻¹, and a sheath gas temperature of 350°C at a flow rate of 11 l min⁻¹. The nebulizer pressure was set to 35 psig (~2.4 bar). The fragmentor voltage was set at 150 V. The spectra were scanned in fragmentation mode (Auto MS/MS) at a range of 95–1000 m/z for both MS and MS/MS. For each cycle, the top 5 precursors were selected for fragmentation using fixed collision energies of 30.00 and 50.00 eV. Active exclusion was enabled for 0.30 min after the acquisition of one spectrum to prevent redundant sampling of high-abundance ions. LC–MS system operation and data acquisition were supervised by Agilent MassHunter software.

The acquired LC–MS data were initially processed using MS-DIAL for peak detection, alignment, and metabolite annotation. The processed data were subsequently imported into MetaboAnalyst for further statistical analysis. Within MetaboAnalyst, missing values were imputed, followed by log transformation and data filtering to improve data quality and ensure normal distribution. The processed dataset was then subjected to multivariate analysis, specifically partial least squares discriminant analysis (PLS-DA), to identify patterns of variation and group separation among the samples. To determine statistically significant metabolites, univariate analysis was performed using one-way ANOVA, followed by Fisher’s post hoc test. Multiple testing correction was applied using the false discovery rate (FDR), and metabolites with an FDR ≤ 0.05 were considered significantly altered.

**Supplemental Figures**

**Figure S1.** **JAL family members are repressed by pathogen-associated signals.** (A) Confocal images showing *pJAL8* reporter activity during infection with *R. pseudosolanacearum* over a 96-hour time course. GFP fluorescence is shown in green. Scale bar, 100 μm. (B) Reporter activity of *pJAL10* and *pJAL20* following flg22 treatment for 3 h. Scale bar, 75 μm. (C) Reporter activity of *pJAL10* and *pJAL20* following *Fusarium oxysporum* infection at 72 hpi. Scale bar, 75 μm. (D) Reporter activity of pJAL20 following *Rhizoctonia solani* infection. Bright-field scale bar, 10 μm; confocal scale bar, 90 μm. Statistical significance was determined using the Kruskal–Wallis test followed by Dunn’s multiple comparison test (*P ≤ 0.05, **P ≤ 0.01, ***P ≤ 0.001). Three independent biological replicates were performed (n = 15 per experiment).

**Figure S2**. **Auxin positively regulates JAL10 and JAL20 expression.** (A) Confocal images showing pJAL10 and pJAL20 reporter activity following treatment with 1 μM IBA. GFP fluorescence is shown in green and cell walls stained with propidium iodide in red. (B) Quantification of fluorescence intensity following IBA treatment. Data represent three independent biological replicates (n = 15 per experiment). (C, D) Relative transcript levels of *JAL10* and *JAL20* in the auxin receptor mutant *tir1* under control and infection conditions. Statistical significance was determined using two-way ANOVA followed by Fisher’s LSD test.

**Figure S3.** **Validation of JAL functional mutants and invasion assay experimental setup**. (A) Semi-quantitative RT–PCR analysis of *JAL10* and *JAL20* expression in Col-0, knockout mutants, and overexpression lines. *ACTIN* was used as an internal control. (B) Schematic representation of the *in vitro* invasion assay. (C) Representative confocal images showing progression of *R. pseudosolanacearum-mCherry* during invasion assays from 24 to 120 hpi in JAL10 and JAL20 functional mutants. Bacterial fluorescence is shown in red and cell walls stained with calcofluor white. Scale bar, 50 μm.

**Figure S4.** Representative images of bacterial colonies formed on BG agar plates following CFU assays in JAL10 and JAL20 functional mutants.

**Figure S5**. **Overview of glycoproteomic analysis.** (A) Schematic workflow of the glycoproteomic analysis. (B) Principal component analysis of glycoproteomic profiles from JAL10 and JAL20 functional mutants. (C) Gene Ontology enrichment analysis of the total glycoproteome, including biological process (BP), molecular function (MF), cellular component (CC), and KEGG pathway enrichment.

**Figure S6.** **Distribution of significantly altered glycoproteins in JAL mutants.** (A, B) Principal component analysis of significantly altered proteins in JAL10 and JAL20 functional mutants.(C-F) Relative abundance of significantly altered proteins under control and infection conditions. (G-J) Distribution of enriched biological processes in JAL10 and JAL20 functional mutants and Col-0 under control and infection conditions.

**Figure S7.** **Functional enrichment analysis of significantly altered proteins.** Gene Ontology enrichment analysis of significantly altered proteins in JAL10 and JAL20 functional mutants showing enriched biological process (BP), molecular function (MF), cellular component (CC), and KEGG pathway categories.

**Figure S8. Clustering analysis of glycoproteins in JAL10 functional mutants.** (A) Expression patterns of significantly altered proteins in JAL10 functional mutants under control and *R. pseudosolanacearum* infection conditions. (B) Heatmap showing K-means clustering of significantly altered proteins relative to Col-0 control conditions.

**Figure S9. Clustering analysis of glycoproteins in JAL20 functional mutants.** (A) Expression trends of significantly altered proteins in JAL20 functional mutants under control and R. pseudosolanacearum infection conditions. (B) Heatmap showing K-means clustering of significantly altered proteins relative to Col-0 control conditions.

**Figure S10. Metabolomic profiling of JAL functional mutants.** (A, B) Principal component analysis and distribution of significantly altered metabolites identified using one-way ANOVA with FDR correction (FDR ≤ 0.05). Analyses were performed using MetaboAnalyst. (C, D) Heatmaps showing the top 10% of metabolites identified by PLS-DA analysis in JAL10 and JAL20 functional mutants.

**Figure S11.** **Identification and functional analysis of tomato JAL orthologs.**

(A) Domain architecture of tomato jacalin-associated lectins identified in *Solanum lycopersicum*. (B-D) Tissue-specific transcript abundance of SlJAL1, SlJAL5, and SlJAL9. (E) Relative transcript abundance of *SlJAL1*, *SlJAL5*, and *SlJAL9* following *R. pseudosolanacearum* infection in tomato varieties with differing susceptibility levels. in the Pusa Ruby- highly susceptible variety, Arka Vikas- moderately susceptible variety, and Utkal Kumari- tolerant variety (F) Phenotype of pTRV plants used as a positive control for VIGS efficiency. (G) Validation of transcript silencing in SlJAL1-, SlJAL5-, and SlJAL9-silenced plants. Statistical significance was determined using Student’s *t*-test (*P ≤ 0.05, **P ≤ 0.01, ***P ≤ 0.001).

**Figure S12.** **JAL family members are repressed by pathogen-associated signals.** (A) Reporter activity of *pJAL10* and *pJAL20* following flg22 treatment for 3 h. Scale bar, 75 μm. (B) Reporter activity of *pJAL10* and *pJAL20* following *Fusarium oxysporum* infection at 72 hpi. Scale bar, 75 μm. (C) Reporter activity of pJAL20 following *Rhizoctonia solani* infection. Bright-field scale bar, 10 μm; confocal scale bar, 90 μm. Statistical significance was determined using the Kruskal–Wallis test followed by Dunn’s multiple comparison test (*P ≤ 0.05, **P ≤ 0.01, ***P ≤ 0.001). Three independent biological replicates were performed (n = 15 per experiment).

**Supplemental Tables**

Table S1.	Cytokinin drench assay statistics

Table S2.	Functional mutants JAL10 invasion assay statistics

Table S3.	Functional mutants JAL20 invasion assay statistics

Table S4.	Functional mutants JAL10 drench assay statistics

Table S5.	Functional mutants JAL20 drench assay statistics

Table S6.	Functional mutants JAL10 Glycoproteome data

Table S7.	Functional mutants JAL20 Glycoproteome data

Table S8.	Final number of Glycopeptides filtered with ± 20ppm mass error and 0.01 FDR

Table S9.	Broad biological classification

Table S10.	Significant proteins in JAL10 functional mutants glycoproteome

Table S11.	Z_score_abundance_JAL10 functional mutants glycoproteome

Table S12.	Significant proteins in JAL20 functional mutants glycoproteome

Table S13.	Z_score_abundance_JAL20 functional mutants glycoproteome

Table S14.	MS-peaks for JAL10 functional mutants

Table S15.	PLSDA for JAL10 functional mutants

Table S16.	Annova_JAL10 functional mutants

Table S17.	MS Peaks for JAL20 functional mutants

Table S18.	PLSDA for JAL20 functional mutants

Table S19.	Annova_JAL20 functional mutants

Table S20.	VIGS drench assay

Table S21.	Primer used in this study
